# HUWE1 stimulates mTORC1 activity by enhancing Rheb interaction with mTORC1 and supports de novo pyrimidine synthesis

**DOI:** 10.1101/2025.04.22.650033

**Authors:** Takayuki Ikeda, Sungki Hong, Shota Yoshida, Abhiram Kunamneni, Swayam Prakash Srivastava, Yao Yao, Venkatesha Basrur, Henry Kuang, Maureen Kachman, Heidi Baum, Arya Joshi, Vinamra Swaroop, Aaron Thurman, Oliviamae Kopasz-Gemmen, Siddharth Kandukuri, Om Khuperkar, FNU Pradeepa, Hideto Yonekura, Jiandie D Lin, Ken Inoki

**Affiliations:** Life Sciences Institute, University of Michigan, 210 Washtenaw Avenue, Ann Arbor, MI, 48109, USA; Department of Biochemistry, Kanazawa Medical University School of Medicine, 1-1 Daigaku, Uchinada, Kahoku-gun, Ishikawa, 920-0293, Japan; Department of Pathology, University of Michigan, Ann Arbor, MI, 48109, USA; Department of Cell & Developmental Biology, University of Michigan Medical School, Ann Arbor, MI, 48109, USA; Biomedical Research Core Facilities, University of Michigan, Ann Arbor, MI, 48109, USA; Department of Molecular and Integrative Physiology, University of Michigan, Ann Arbor, MI, 48109, USA; Department of Internal Medicine, University of Michigan, Medical School, 210 Washtenaw Avenue, Ann Arbor, MI, 48109, USA

## Abstract

mTORC1 is a central regulator of cell growth that is directly activated by the small GTPase Rheb on the lysosomal membrane in response to growth factors. We recently reported that polyubiquitinated Rheb facilitates its interaction with mTORC1, leading to mTORC1 activation. However, the molecular mechanisms underlying the ubiquitination of Rheb and its interaction with mTORC1 for activation are not fully understood. In this study, we demonstrate that HUWE1, an E3 ubiquitin ligase, preferentially interacts with ubiquitinated Rheb and is essential for the interaction between Rheb and mTOR, as well as for mTORC1 activation. Additionally, HUWE1 is necessary for Rheb to interact with CAD, a crucial enzyme involved in de novo pyrimidine biosynthesis, and for its activation through the mTORC1-S6K1 pathway. Knocking down HUWE1 in cultured cells or in the liver tissues of mice inhibits mTORC1 activity without affecting the phosphorylation of Akt or TSC2, nor does it affect the lysosomal localization of TSC2 or mTORC1. Furthermore, HUWE1 specifically enhances the expression of CAD without influencing other enzymes involved in de novo pyrimidine synthesis and maintains UMP levels in the hepatocytes of liver tissues. These findings indicate that HUWE1 serves as a key organizer of the Rheb ubiquitin complex, amplifying mTORC1 activity and playing a vital role in stimulating de novo pyrimidine synthesis by enhancing CAD expression and activity.

**Highlights:**

1. HUWE1 and CAD preferentially interact with ubiquitinated Rheb.
2. HUWE1 further ubiquitinates Rheb and supports Rheb interaction with mTORC1 and CAD.
3. HUWE1 stimulates mTORC1 activity through its E3 ligase activity.
4. HUWE1 also supports pyrimidine synthesis by enhancing CAD expression.

## Introduction

The mechanistic target of rapamycin complex 1 (mTORC1) plays a central role in stimulating cellular anabolic processes that are essential for cell growth and proliferation, including protein, lipid, and nucleotide biogenesis ^1^. In response to amino acids and growth factors, mTORC1 is primarily activated by two distinct lysosomal Ras-related small GTPases, Rags and Rheb, on the lysosomal membrane ^2^. While Rags recruit mTORC1 to the lysosomal membrane in response to amino acid availability, growth factors activate Rheb, which directly interacts with and stimulates mTORC1 on the lysosome ^3,4^. A series of recent studies have uncovered how amino acids stimulate Rags by regulating their GTPase activating proteins (GAPs, e.g. GATOR1 for RagA/B, FLCN-FLIPs/LRS for RagC/D) ^5–7^ and guanine-nucleotide exchange factors (GEFs, e.g., SLC38A9 for RagA/B and Ragulator for RagC/D) ^8^. For Rheb activation, Akt activated by the growth factor directly phosphorylates and inhibits the function of TSC2, the GAP for Rheb, by dislocating the TSC complex away from the lysosomal membrane, thereby providing a permissive condition to Rheb for its GTP-loading and activation ^9–18^. In addition, recent studies demonstrated that the lysosomal localization of TSC2 is also regulated by amino acids where the inactive Rags tether the TSC complex to the lysosome ^19^. However, GEFs for Rheb, which convert GDP-bound inactive Rheb to GTP-bound active Rheb, have not been convincingly demonstrated.

Furthermore, recent our studies demonstrated that polyubiquitinated Rheb induced by amino acids facilitates Rheb-mTORC1 interaction and subsequent mTORC1 activation ^20,21^. Similar to the previously demonstrated mechanism underlying amino acid-dependent interaction between Rags and TSC2, we found that Ataxin 3 (ATXN3), the Rheb deubiquitinase, displays a strong binding preference for the inactive Rag heterodimer and translocates to the lysosome where it deubiquitinates Rheb under amino acid starved conditions. Thus, the antiparallel spatial movement of mTORC1 and ATXN3 or the TSC complex between cytosol and lysosomal surface ensures mTORC1 is effectively activated or inactivated in response to both growth factor and amino acid availability. However, it has not been well understood what ubiquitin ligases contribute to Rheb polyubiquitination and its modifications and support mTORC1 activation.

HUWE1, also known as an ARF-binding protein or Mule, is a large E6AP-type E3 ubiquitin ligase containing multiple protein-protein interaction domains ^22^. HUWE1 is known to ubiquitinate multiple lysine residues (e.g., K6, K48, and K63) to generate either poly-ubiquitin chains or branches on target proteins, leading to protein degradation or stabilization, respectively ^23,24^. Known HUWE1-mediated degradation targets include both oncogenic proteins (e.g., N-Myc, c-Myc, and Mcl-1) and a tumor suppressor protein (p53), indicating that HUWE1 functions in both pro-apoptotic and anti-apoptotic pathways ^25–29^. HUWE1 also inhibits autophagy induction by degrading WD Repeat Domain Phosphoinositide-Interacting Protein 2 (WIPI2), a key protein promoting autophagosome formation ^30^. In contrast to its role in protein degradation, HUWE1 can generate K63-linked ubiquitin on c-Myc for its activation and K48 branches to K63 ubiquitin chains of TNF receptor-associated factor 6 (TRAF6) and amplify NF-κB signaling ^24,25^. Recent studies also demonstrated that HUWE1 also functions as an E3 ligase for NEDD8, a ubiquitin-like protein, which protects ubiquitinated proteins from proteasome-mediated degradation ^31^.

The carbamoyl-phosphate synthetase 2, aspartate transcarbamoylase, dihydroorotase (CAD) is a multifunctional enzyme that catalyzes the first three steps of de novo pyrimidine synthesis ^32^. The enzyme activity of CAD is stimulated by S6K1-dependent phosphorylation, indicating that CAD is a key substrate of the mTORC1-S6K1 pathway ^33,34^. A recent study also reported that Rheb and RhebL1 (Rheb2), which is predominantly expressed in the brain, interact with CAD and stimulate its activity independent of mTORC1 activity ^35^. RhebL1 shows a much higher affinity for CAD compared to Rheb. Furthermore, co-localization of CAD with RhebL1 on the lysosome membrane is diminished by a farnesyltransferase inhibitor that blocks membrane localization of RhebL1, suggesting that RhebL1 plays an important role in the lysosomal localization of CAD. Furthermore, inhibition of the second and third enzymes in the de novo pyrimidine pathway, DHODH and UMPS, respectively, shows little effect on cellular mTORC1 activity ^36,37^. These observations suggest that CAD is a downstream effector of RhebL1 and Rheb and a substrate of the mTORC1-S6K1 pathway.

In this report, we identified that HUWE1 functions as a positive regulator for mTORC1 activation upstream of Rheb, a direct activator of mTORC1. The ablation of HUWE1 mitigates amino acid- or growth factor-induced mTORC1 activation without affecting mTORC1 or TSC2 lysosomal localization and phosphorylation status of Akt and TSC2. Importantly, the ablation of HUWE1 diminishes the interaction of Rheb with mTOR and CAD. Furthermore, we found that HUWE1 also plays a key role in enhancing CAD transcription. Consistent with these observations in cultured cells, in the liver tissues lacking HUWE1 in their hepatocytes, mTORC1 activity, and CAD expression are significantly reduced, resulting in the reduction de novo pyrimidine synthesis pathway. Taken together, we propose that the HUWE1 serves as not only a key organizer of the Rheb-ubiquitin complex, which facilitates Rheb-dependent mTORC1/CAD activation, but also an upstream of a positive regulator of CAD transcription, thus amplifying de novo pyrimidine synthesis.

## Results

### Ubiquitinated Rheb displays a strong binding preference for HUWE1 and CAD

To identify Rheb ubiquitin ligases and possible Rheb GEFs, which have a strong binding preference for nucleotide-free small GTPases in general, we generated a nucleotide-free Rheb mutant in which the conserved glycine in the phosphate-binding loop (P-loop) was replaced with an alanine (Fig. 1A). This Rheb G18A mutant is predicted to lose guanine nucleotide binding and forms a nucleotide-free form of Rheb (Fig. 1A). An equivalent mutation in the P-loop of Ras (G15A) abrogates guanine-nucleotide binding and promotes strong and stable interaction with its activator, cdc25, a GEF for Ras ^38^. The nucleotide-binding status of the Rheb G18A mutant and other previously reported Rheb mutants (S16H: GTP form, D60I: GDP form, D60K: nucleotide-free form) was examined under ^32^P orthophosphate labeling conditions ^39–42^. As expected, the G18A Rheb mutant failed to bind guanine nucleotides, while the S16H and D60I mutants exclusively bound to GTP and GDP, respectively (Fig. 1B). The D60K mutant largely diminished nucleotide binding with a small remaining GDP (Fig. 1B). Simultaneously, we also monitored levels of ubiquitination in wild-type, S16H, D60I, and G18A. We observed that while all the Rhebs generated their polyubiquitinated forms migrated above 250 kD, the Rheb D60I and G18A mutants displayed much higher levels of ubiquitination compared to those in wild-type and the S16H mutant (Fig. 1C). The blockade of membrane tethering of Rheb by introducing C181S mutation on the Rheb G18A mutant reduced levels of ubiquitination, suggesting that membrane localization play a role in supporting the ubiquitination of the Rheb G18A mutant (Fig. 1C). Thus, nucleotide-free Rheb G18A mutant would act as a potential bait to capture possible Rheb GEF proteins and/or ubiquitin ligases that ubiquitinate Rheb on the membrane. To isolate such Rheb regulators, we performed protein purification using the Rheb G18A mutant (nucleotide-free form with high ubiquitination) and Rheb S16H mutant (GTP-bound form with low ubiquitination) as baits (Fig. 1D). Silver staining demonstrated that two specific bands were predominantly co-immunoprecipitated (co-IPed) with the Rheb G18A mutant. Through mass spectrometry analyses, we identified that these bands are CAD and HUWE1. Among these Rheb mutants, G18A and D60I Rheb, which are highly ubiquitinated (Fig. 1C), displayed a strong binding preference for endogenous HUWE1 and CAD (Fig. 1E). Furthermore, as the blockade of membrane localization of Rheb reduced its ubiquitination (Fig. 1C), levels of co-IPed endogenous HUWE1 and CAD with the Rheb G18A/C181S mutant were largely decreased comparing to those with the Rheb G18A mutant (Fig. 1F). These observations suggest that rather than nucleotide-binding status of Rheb, levels of ubiquitination or membrane localization may play a role in enhancing the binding of Rheb to HUWE1 and CAD. To dissect further this binding module under more physiological conditions, we tested the interaction of endogenous Rheb with HUWE1 and CAD in growing HEK293T and HeLa cells. While endogenous Rheb could co-IP a very small amount of endogenous HUWE1 and CAD, levels of co-IPed HUWE1 and CAD with endogenous Rheb were significantly increased when N-ethylmaleimide (NEM), a cysteine peptidase and deubiquitinase inhibitor, was added during cell lysis and immunoprecipitation (Fig. 1G). These results suggest that the preservation of the ubiquitinated form of Rheb maintains its binding affinity to both HUWE1 and CAD.

**Figure 1.**
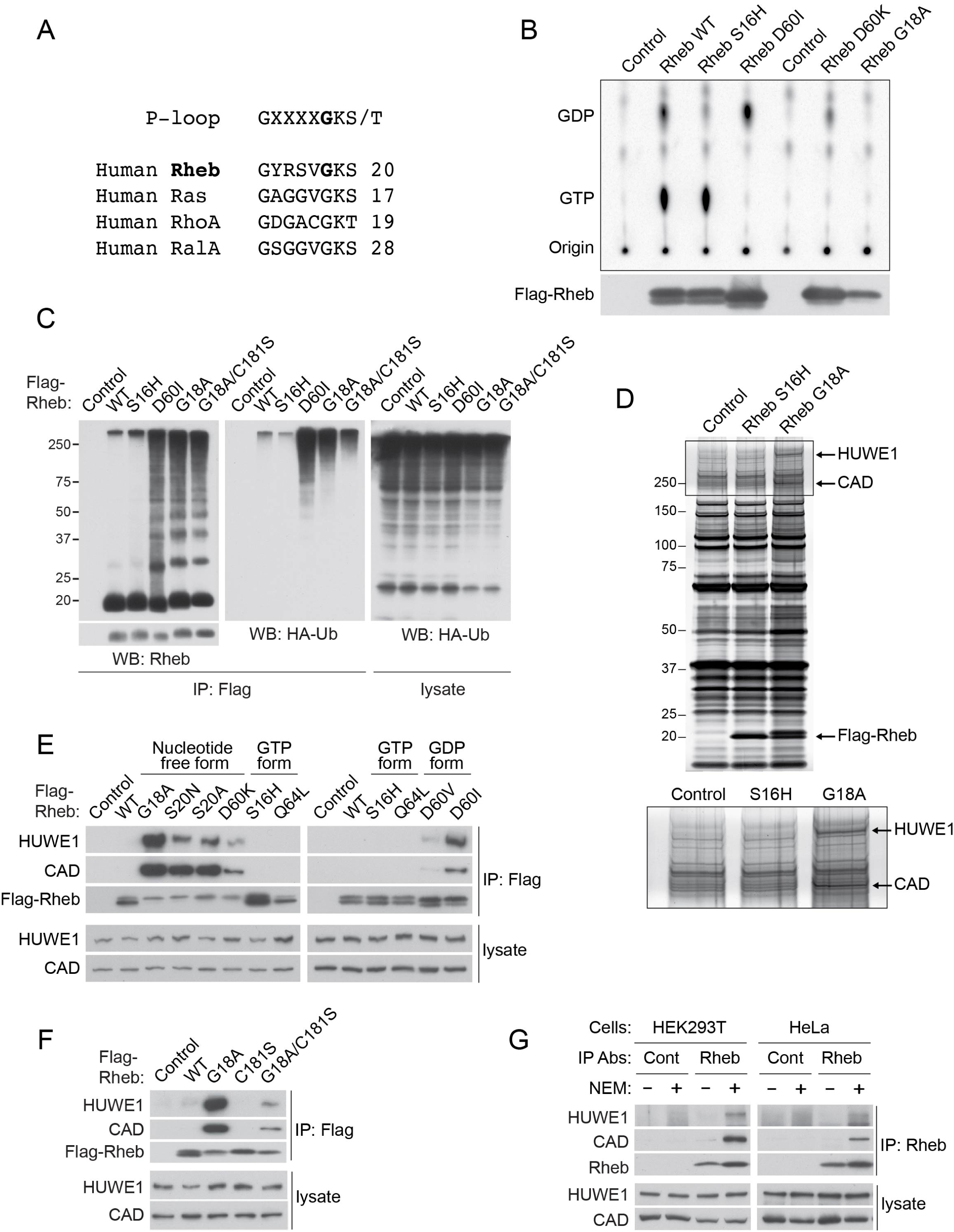
Ubiquitinated nucleotide-free or GDP-bound form of Rheb preferentially interacts with endogenous HUWE1 and CAD. **(A) The GXXXXGKS/T domain, named P-loop, is conserved in the Ras family of small GTPases.** Rheb has a conserved Gly18 residue (bold). **(B) Nucleotide binding status of Rheb mutants.** HEK293T cells transiently expressing the indicated Rheb mutants were labeled with ^32^P orthophosphate. Levels of GTP and GDP bound to the IPed Flag-Rheb were shown. **(C) Nucleotide-free and GDP-bound forms of Rheb mutants are highly ubiquitinated in a manner dependent on the membrane.** HEK293T cells were co-transfected with HA-Ub, and the indicated Flag-Rhebs (wild-type, S16H (GTP-bound form), D60I (GDP-bound form), G18A (nucleotide-free form), or G18A/C181S (nucleotide-free/membrane unbound) mutants). Flag-Rheb was IPed with Flag antibody-conjugated beads (Flag-beads), and levels of high-molecular-weight Rhebs were monitored by Rheb or HA antibody. **(D) Highly ubiquitinated nucleotide-free Rheb G18A mutant strongly interacts with endogenous HUWE1 and CAD.** HEK293T cells were transfected with Flag-Rheb S16H (GTP-bound form) or G18A (nucleotide-free form) mutant. Flag-Rheb was IPed with Flag-beads, and co-IPed proteins were eluted and visualized by silver staining and followed by mass spectrometry analyses. **(E) Nucleotide-free or GDP-bound form of Rheb preferentially interacts with endogenous HUWE1 and CAD.** HEK293T cells transiently expressing the indicated Rhebs were subjected to IP with Flag-beads. Levels of co-IPed endogenous HUWE1 and CAD were monitored by western blotting. **(F) Blocking membrane tethering of Rheb G18A mutant mitigates its interaction with HUWE1 and CAD.** C181S mutation was introduced in wild-type Rheb and G18A Rheb mutant. The experiments were performed as 1E. **(G) Inhibition of deubiquitinase activity increases interaction between endogenous Rheb and HUWE1 or CAD.** Endogenous Rheb was subjected to IP, and levels of co-IPed endogenous HUWE1 and CAD were monitored in the presence or absence of NEM, a deubiquitinase inhibitor, in the lysates.

We also examined the subcellular localization of endogenous Rheb, HUWE1, and CAD by performing subcellular fractionation and immunofluorescence staining assays. Similar to TSC2, HUWE1 and CAD are primarily expressed in the cytoplasmic and membrane fractions (Fig. S1A) and partially co-localize with Rheb in the perinuclear region (Fig. S1B, S1C, S1D, S1E, and S1F).

### HUWE1 is required for Rheb to interact with CAD and mTOR

To understand the nature of HUWE1 and CAD interactions with Rheb, we first asked if HUWE1 interacts with CAD. HUWE1 co-IPed CAD, and this interaction was modestly enhanced by growth factor/amino acid stimulation (Fig. 2A). This result suggests that HUWE1 and CAD may be in the same complex with Rheb. To determine the key domains of CAD or HUWE1 that interact with Rheb, we generated various truncation mutants of CAD and HUWE1. Co-IP experiments demonstrated that the Rheb G18A mutant effectively co-IPed CAD that contains the CPSase B domains (Fig. 2B), which is consistent with the previous observations ^35^. On the other hand, the HECT domain, but not the UBA (ubiquitin-associated) domain, of HUWE1 effectively co-IPed Rheb G18A mutant (Fig. 2C). The interaction of HUWE1 with Rheb G18A did not require HUWE1’s catalytic activity as the catalytic inactive HUWE1 C4341S mutant or the HECT domain with a C4341S mutation displayed an equivalent affinity to their wild-type proteins to interact with Rheb (Fig. 2C). Interestingly, both full-length and the HECT domain of HUWE1 effectively co-IPed the CPSase domains of CAD (Fig. 2D and 2E). These data suggest that the HECT domain of HUWE1 or the CPSase domain of CAD may play a key role in the association between CAD and Rheb or HUWE1 and Rheb, respectively (Fig. 2F).

**Figure 2.**
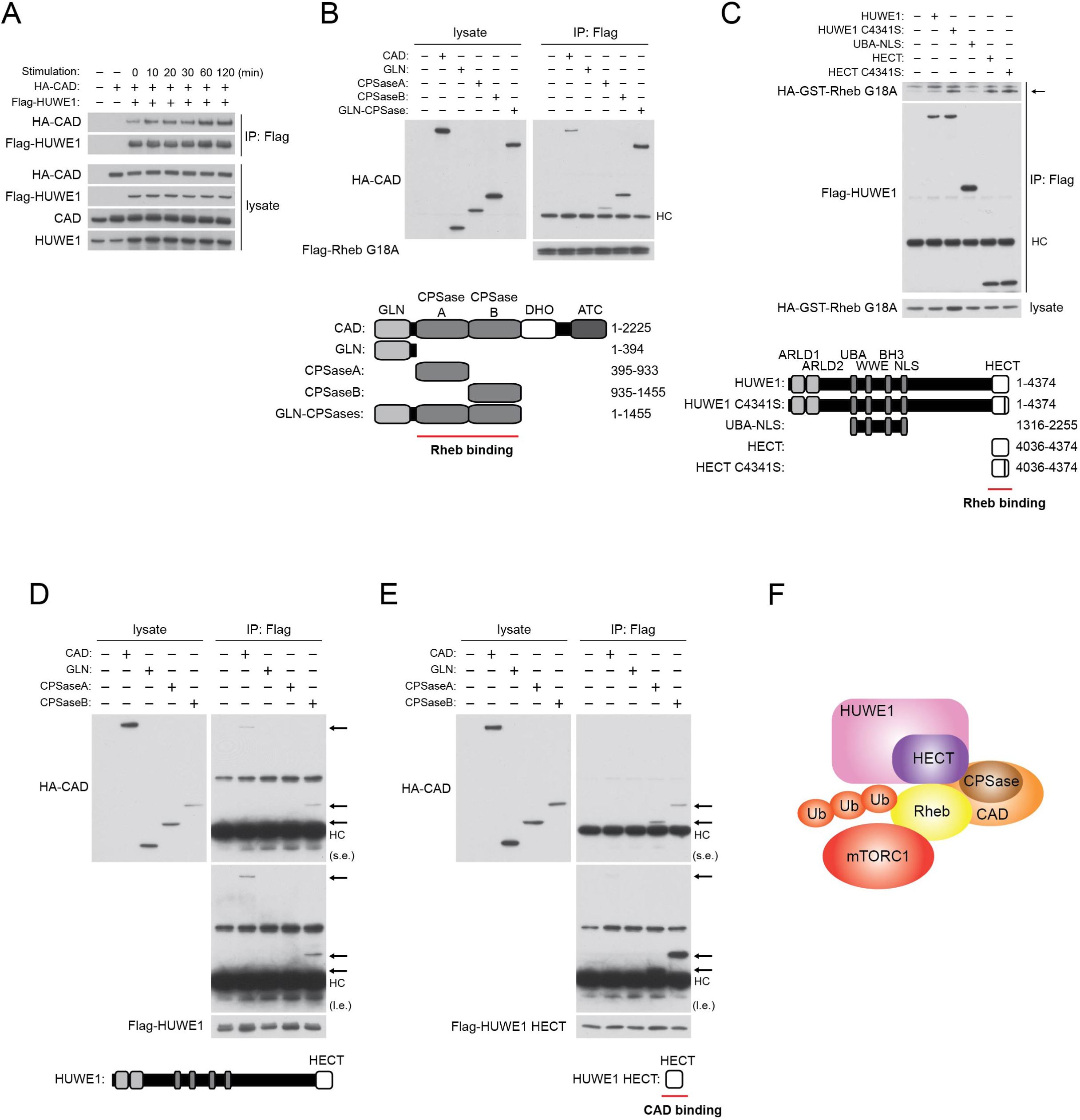
Characterization of binding modules of Rheb-HUWE1-CAD interaction. **(A) HUWE1 interacts with CAD.** HEK293T cells transiently expressing HA-CAD and Flag-HUWE1 at low levels were starved with HBSS and then stimulated with DMEM containing 10 % FBS. Flag-HUWE1 was IPed, and levels of co-IPed HA-CAD were monitored. Total CAD and HUWE1 in the lysate were also determined. **(B) The Rheb G18A mutant interacts with the CPSase domain of CAD.** HEK293T cells expressing Flag-Rheb G18A and the indicated HA-CAD and HA-CAD fragments were lysed and IPed with Flag-beads. Co-IPed HA-CAD was monitored. HC denotes IgG heavy chain. **(C) The Rheb G18A mutant interacts with the HECT domain of HUWE1.** HEK293T cells expressing the indicated Flag-HUWE1 and Flag-HUWE1 fragments and HA-GST-Rheb G18A were lysed and IPed with Flag-beads. Co-IPed HA-GST-Rheb G18A was monitored. HC denotes IgG heavy chain. The arrow indicates co-IPed Rheb. **(D, E) HUWE1 and its HECT domain interact with the CPSase domain of CAD.** HEK293T cells expressing Flag-HUWE1 (D) or Flag-HUWE1-HECT (E) with the indicated HA-CAD and HA-CAD fragments were lysed and IPed with Flag-beads. Co-IPed HA-CAD was monitored. HC denotes IgG heavy chain. The arrow indicates co-IPed CAD. s.e. and l.e. indicate short and long exposure, respectively. **(F) Hypothetical binding model of HUWE1-CAD-Rheb interaction.** The HECT domain of HUWE1, which can recognize its substrates and ubiquitinated residues, and the CPSase domain of CAD are required for the interactions between HUWE1, CAD, and Rheb.

To determine if CAD is required for Rheb interaction with HUWE1, or HUWE1 is required for Rheb interaction with CAD, we depleted either CAD or HUWE1 by shRNA-mediated knockdown (KD) and monitored Rheb interactions with the other. While CAD KD had little effect on the interaction between Rheb G18A and HUWE1, HUWE1 KD largely diminished Rheb G18A interaction with CAD, indicating that HUWE1 is required for CAD to interact with Rheb G18A mutant (Fig. 3A). Importantly, HUWE1 KD but not CAD KD also disrupted the interaction between Rheb G18A mutant and mTOR, suggesting that HUWE1 may play a key role in the association between Rheb and mTOR (Fig. 3A). To investigate whether HUWE1’s role in the enhancing Rheb interaction with mTOR is cell-type specific, CAD or HUWE1 was ablated in MEFs, and the effect of CAD KD or HUWE1 KD on the interaction between endogenous Rheb and mTOR was monitored. Similar to the observations in HEK293T cells using Rheb G18A, CAD KD demonstrated little effect on growth factor/amino acid-induced interaction between endogenous Rheb and mTOR (Fig. 3B). Importantly, HUWE1 KD largely inhibited growth factor/amino acid-induced endogenous Rheb interaction with mTOR (Fig. 3B), indicating that HUWE1 plays a key role in enhancing the interaction between Rheb and mTORC1 as well as CAD.

**Figure 3.**
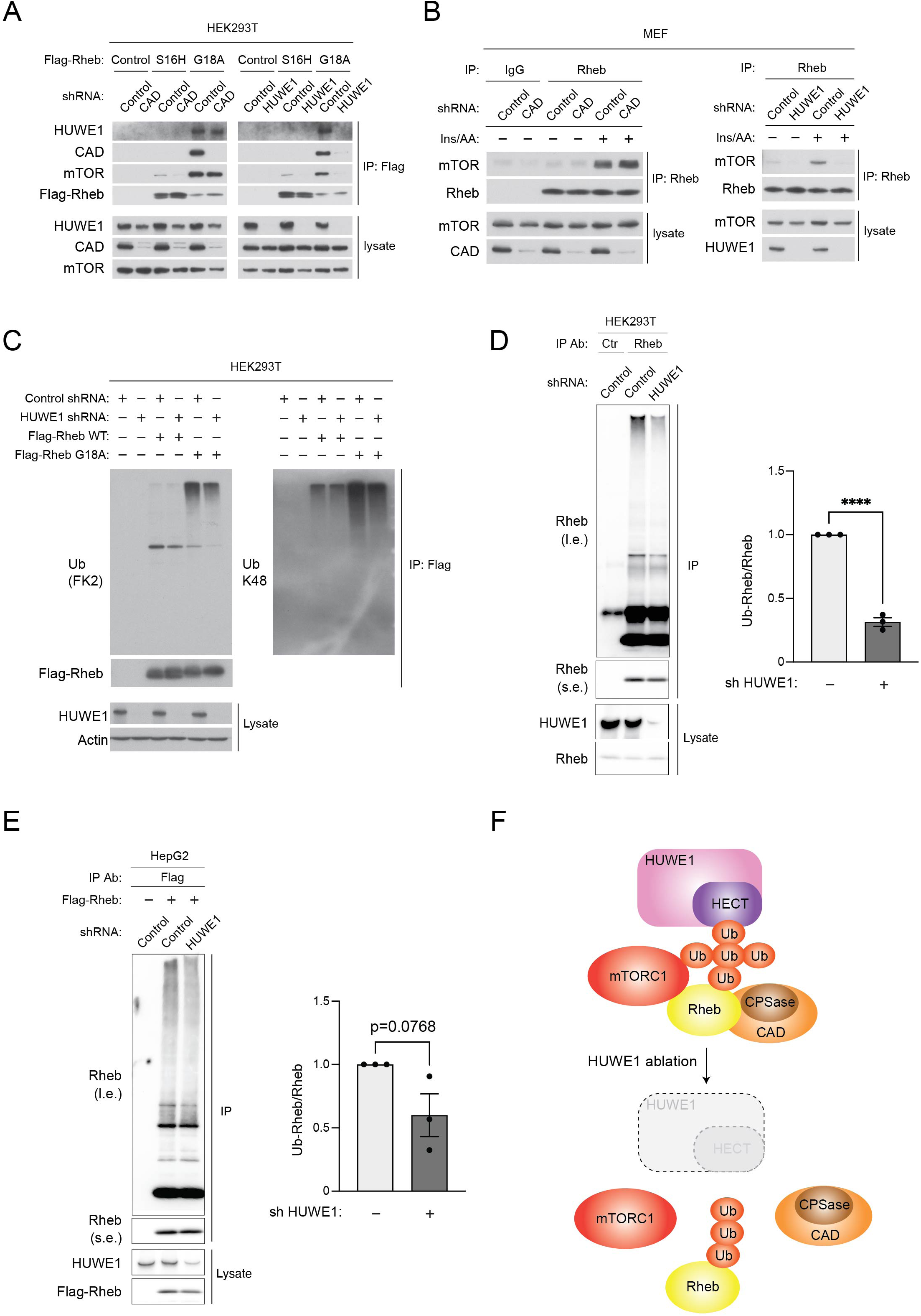
HUWE1 is required for Rheb to interact with CAD and mTOR. **(A) HUWE1 is required for the Rheb G18A mutant to interact with CAD and mTOR.** HEK293T cells expressing control, HUWE1, or CAD shRNA were transfected with the Flag-Rheb S16H (GTP form) or Flag-Rheb G18A (nucleotide-free form) mutant. Cells were lysed and Flag-Rhebs were IPed with Flag-beads. Levels of co-IPed endogenous HUWE1, CAD, and mTOR were monitored. **(B) HUWE1, but not CAD, is required for endogenous Rheb to interact with mTOR.** Endogenous Rheb was IPed from MEFs expressing control, CAD, or HUWE1 shRNA under in vivo crosslinking conditions. Levels of co-IPed endogenous mTOR were monitored. **(C) Ablation of HUWE1 decreases Rheb ubiquitination**. HUWE1 is knocked down with the shRNA in HEK293T cells expressing Flag-tagged wild-type Rheb and G18A Rheb mutant. Flag-Rhebs were IPed, and levels of ubiquitinated Rheb were monitored by pan ubiquitin or K48-linked ubiquitin antibody. **(D and E) Ablation of HUWE1 decreases endogenous Rheb ubiquitination.** HUWE1 is knocked down with the shRNA in HEK293T cells (D) and HepG2 cells (E). Endogenous Rheb was IPed, levels of ubiquitinated high-molecular weight Rheb (Ub-Rheb) were monitored. Quantification was displayed as a ratio of Ub-Rheb/non-Ub-Rheb. ****p<0.0001, mean±SEM, n=3 (D) and p=0.0768, mean±SEM, n=3 (E). Note that the KD efficiency of HUWE1 in HepG2 cells was lower than that in HEK293T cells. **(F) Hypothetical model of HUWE1 function in forming Ub-Rheb-mTORC1-CAD complex.** HUWE1 plays a critical role in enhancing Rheb to interact with both mTORC1 and CAD possibly through Rheb ubiquitin modification.

### HUWE1 regulates Rheb ubiquitination

HUWE1 preferentially interacts with nucleotide-free or GDP-bound forms of Rheb mutants, which are highly ubiquitinated in the cells (Fig. 1B, C, and E). In addition, HUWE1 plays a role in enhancing Rheb binding to mTOR. These observations suggest that HUWE1 acts as a key ubiquitin ligase that supports Rheb interaction with mTOR or possibly functions as a GEF-like protein. We examine if HUWE1 displays any GEF-like activity toward Rheb using purified HUWE1 and Rheb protein in vitro and found that HUWE1 did not display a detectable guanine-nucleotide exchange activity for Rheb (Fig. S2A).

Previous studies demonstrated several distinct ubiquitin ligases, including Siah1 ^43^, RNF152 ^44^, or DCAF1 ^45^, ubiquitinate and degrade Rheb in response to nitric oxide, growth factor starvation, and glucose scarcity, respectively. Interestingly, we observed that while the polyubiquitinated Rheb (Ub-Rheb) is gradually subjected to its proteasome and lysosomal degradation, Ub-Rheb forms a hetero-multimer with non-ubiquitinated Rheb and facilitates their interaction with mTORC1 on the lysosome and its subsequent activation ^20^. We recently identified that the CUL3-Rbx1-KLHL9 complex ubiquitinates Rheb to form “active” Ub-Rheb for mTORC1 activation ^21^, whereas ATXN3 deubiquitinates Rheb and inhibits mTORC1 activation under amino acid starved conditions ^20^. To investigate if HUWE1 affects levels of Rheb ubiquitination, the effect of HUWE1 KD on wild-type and G18A Rheb ubiquitination was monitored in HEK293T. Under Rheb overexpressed conditions, while the effect was moderate, levels of total or K48-linked ubiquitination of both wild-type and G18A Rheb were consistently reduced in HUWE1 KD HEK293T cells (Fig. 3C). Similarly, levels of polyubiquitinated high-molecular weight endogenous Rheb were also reduced in HUWE1 KD HEK293T cells (Fig. 3D). The reduction of Ub-Rheb was also observed in HepG2 cells (Fig. 3E). Since the ablation of HUWE1 did not eliminate Ub-Rheb, these observations suggest that HUWE1 is not a primary E3 ubiquitin ligase to generate main polyubiquitinated form of Rheb, but it may act as a key ubiquitin modifier working with other ubiquitin ligases. This idea is also supported by the fact that HUWE1 showed a stronger binding preference for the Rheb mutants, which are highly ubiquitinated (Fig. 1B, C, and E), or endogenous Rheb, whose deubiquitination was inhibited (Fig. 1G). Together, these observations suggest that HUWE1 may play a key role in the interaction between Rheb and mTOR or CAD through its ubiquitin ligase activity, likely by modifying (e.g., branching) the ubiquitinated form of Rheb (Fig. 3F).

### HUWE1 plays an important role in growth factor/amino acid-induced mTORC1 activation

Given the important role of HUWE1 in stimulating the interaction between Rheb and mTOR, we tested the effect of HUWE1 ablation on cellular mTORC1 and mTORC2 activity. Importantly, HUWE1 KD showed a substantial decrease in mTORC1 activity under normal cell culture conditions (basal) and largely attenuated mTORC1 activation in response to growth factor/amino acid stimulation, as determined by levels of S6K1 and 4EBP1 phosphorylation (Fig. 4A). Similar observations were also seen in HEK293T and HeLa cells (Fig. S2B and S2C). Time course studies also demonstrated that the activation of mTORC1 was largely delayed and mitigated in HUWE1 KD cells (Fig. 4B). While initial induction of Akt, TSC2, or ERK phosphorylation was slightly reduced 5 min after growth factor/amino acid stimulation in HUWE1 KD cells, they quickly recovered to similar or higher levels of phosphorylation in later time points compared to those in control cells (Fig. 4B). Growth factors stimulation, such as insulin treatment leads to the dissociation of TSC2 from the lysosome membrane in MEFs ^46^. Consistent with the little effect of HUWE1 KD on TSC2 phosphorylation (Fig. 4B), insulin treatment sufficiently induced TSC2 dissociation from the lysosome in HUWE1 KD MEFs as in wild-type MEFs (Fig. S2D). It is well known that HUWE1 ubiquitinates and degrades p53, a tumor suppressor that negatively regulates mTORC1 activity ^26,47^. However, similar to the effects in wild-type MEFs, HUWE1 KD significantly inhibited growth factor-induced mTORC1 activity without influencing Akt phosphorylation in MEFs derived from p53 KO embryos (Fig. 4C) ^48^. These observations suggest that HUWE1 plays an important role in regulating mTORC1 activity independent of p53 function and by acting downstream of Akt and TSC2, which is consistent with the observations that HUWE1 acts at the point of Rheb interaction with mTORC1 (Fig. 3A and 3B). If HUWE1 supports Rheb to interact with mTORC1 for its activation, amino acid-induced mTORC1 activation should also be inhibited in HUWE1 KD cells without affecting lysosomal mTORC1 localization. As expected, HUWE1 KD also inhibited leucine-induced mTORC1 activation in MEFs (Fig. 4D) without affecting amino acid-induced lysosomal mTOR localization (Fig. 4E). Intriguingly, HUWE1 KD slightly but consistently decreased CAD expression (Fig. 4A, 4B, and 4D). To investigate if the ligase activity of HUWE1 plays an important role in the activation of mTORC1, we transiently introduced wild type or the catalytic inactive HUWE1 C4341S mutant with HA-S6K1 as a reporter in HUWE1 KD HEK293T cells. While the C4341S mutation in HUWE1 or its HECT domain did not compromise their binding affinity toward Rheb (Fig. 2C), the reduction of S6K1 phosphorylation in the HUWE1 KD cells was restored by re-expressing wild-type HUWE1 but not by the HUWE1 C4341S mutant (Fig. 4F). Increasing CAD expression in HUWE1 KD cells did not influence the effects of wild-type HUWE1 or HUWE1 C4341S mutant on mTORC1 activity, indicating that the E3 ubiquitin ligase activity of HUWE1 contributes to the mechanism underlying HUWE1-dependent mTORC1 activation.

**Figure 4.**
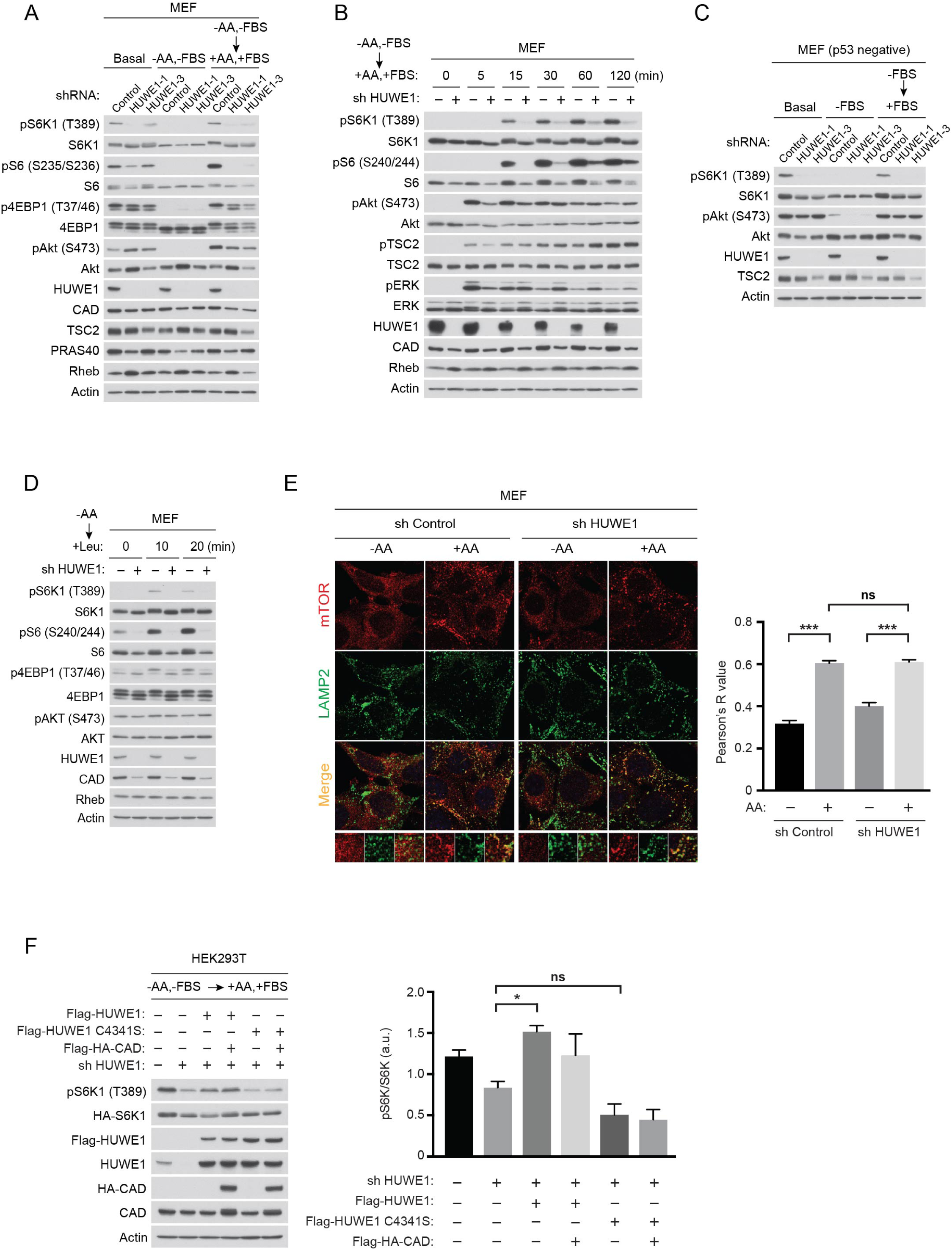
HUWE1 enhances mTORC1 activity. **(A) HUWE1 is required for growth factor/amino acid-induced mTORC1 activation.** MEFs were infected with control (scramble) or two distinct HUWE1 shRNAs, and levels of mTORC1 and mTORC2 activity and the indicated protein expression were monitored under steady-state culture (Basal), HBSS starved (1 hr, HBSS), or growth factor/amino acid-stimulated (20 min, DMEM containing 10% FBS) conditions. **(B) Knockdown of HUWE1 delays and reduces growth factor/amino acid-induced mTORC1 activation (time course).** MEFs expressing HUWE1 shRNA were starved with HBSS for 60 min and then stimulated with DMEM containing 10% FBS for the indicated times. Levels of mTORC1, mTORC2, and the indicated protein expression were monitored. **(C) HUWE1 promotes growth factor-induced mTORC1 activation independent of p53.** MEFs derived from p53 null embryo were infected with control (scramble) or two distinct HUWE1 shRNAs, levels of mTORC1 and mTORC2 activity were monitored under steady-state culture (Basal), growth factor-starved (1 hr, FBS-free medium), or growth factor-stimulated (20 min, 10% FBS) conditions. **(D) HUWE1 is required for amino acid-induced mTORC1 activation.** MEFs were infected with HUWE1 shRNA, and levels of mTORC1 and mTORC2 activity and the indicated protein expression were monitored under leucine-starved (50 min) or leucine-stimulated (10 or 20 min) conditions. **(E) HUWE1 is dispensable for amino acid-induced lysosomal mTOR localization.** MEFs expressing control or HUWE1 shRNA were amino acid-starved for 50 min and then stimulated with 1x amino acids for 10 min. Amino acid-induced lysosomal mTOR localization was monitored by performing double immunofluorescence staining with mTOR (red) and LAMP2 (green) antibodies. Pearson’s correlation coefficient, R values, were determined for quantifying co-localization of mTOR and LAMP2 and graphed as a ***p<0.001, ns (not significant), mean±SEM, n=28 (sh Control, -AA), n=38 (sh Control, +AA), n=23 (sh HUWE1, -AA), n=29 (sh HUWE1, +AA). **(F) The E3 ligase activity of HUWE1 is required for mTORC1 activation.** HEK293T cells ablated endogenous HUWE1 were transiently transfected with Flag-HUWE1 or the catalytically inactive Flag-HUWE1 C4341S mutant in the presence or absence of HA-CAD with HA-S6K1 as a reporter. Cells were starved for 60 min and stimulated with DMEM containing 10% FBS for 60 min. Levels of S6K1 phosphorylation and the indicated proteins were monitored (left panels). Levels of pS6K1/HA-S6K1 were quantified (right graph). Data were expressed as a *p<0.05, ns (not significant), mean±SEM, n=3.

### HUWE1 plays a key role in stimulating mTORC1 activation in vivo

To test the role of HUWE1 in the regulation of mTORC1 activity in a more physiologically relevant setting, hepatocyte-specific HUWE1 KO mice were generated by crossing female *Huwe1 flox/flox* (*Huwe1^f/f^*) mice with male *Albumin-Cre* (*Alb-Cre*) mice. The *Huwe1 ^f/y^*male mice (*Huwe1* is an X-linked gene) as well as *Huwe1^f/f^*female mice expressing *Alb-Cre* (*Huwe1 LKO* mice) grew normally and lived at least two years as wild-type mice (unpublished observations). Consistent with the observation in the cultured cells, 12-week-old male *Huwe1 LKO* mice fed with normal chaw displayed significantly reduced mTORC1 activity monitored by levels of phosphorylated S6 (S240/244), the sites directly phosphorylated by S6K1 in their liver tissues ^49,50^ (Fig. 5A) without affecting Akt phosphorylation (S479) and hepatocyte nucleolus number (Fig. S3A), supporting the idea that HUWE1 specifically enhances mTORC1 activity downstream of Akt. Similar to the results of immunostaining, immunoblot analyses also demonstrated that levels of S6K1 and 4EBP1 phosphorylation, two mTORC1 substrates, and S6 phosphorylation were significantly decreased in the liver tissue lysates from *Huwe1 LKO* mice (*Huwe1^-/y^)* compared to those from control mice (*Huwe1^f/y^*) (Fig. 5B).

**Figure 5.**
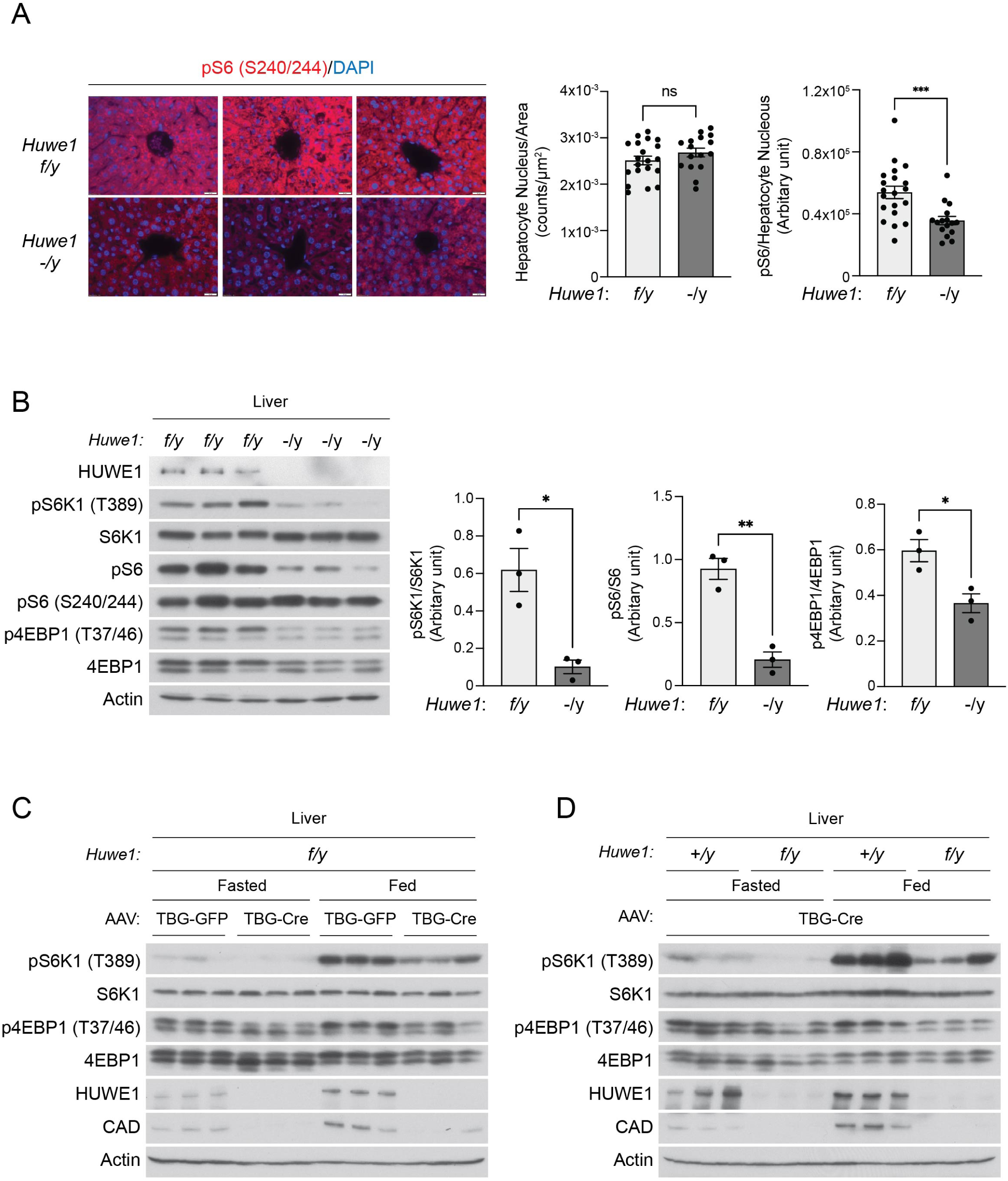
HUWE1 supports mTORC1 activity in the murine liver tissues. **(A and B) Congenital ablation of HUWE1 in hepatocytes reduced mTORC1 activity in the murine liver tissues.** Male *Huwe1^flox/y^* and *Huwe1 LKO* (*Huwe1^flox/y^*, Alb-Cre) mice (12-week-old) were starved for 16 hr and fed for 5 hr. mTORC1 activity and nucleus number of liver tissues (right posterior segment VI) were monitored by staining with pS6 (S240/S244) antibody and DAPI. Quantification was demonstrated as the nucleus number of hepatocyte/area (counts/µm^2^) and the intensity of pS6/nucleus number of hepatocyte (arbitrary unit). ***p<0.001, ns (not significant), mean±SEM, n=16∼20 images (A). Levels of mTORC1 activity were determined by western blotting using pS6K1 (T389), p4EBP1 (pT37/46), and pS6 (S240/S244) antibodies. Quantification was demonstrated as a ratio of the phosphorylated protein/total protein, *p<0.05, **p<0.01, mean±SEM, n=3 (B). **(C and D) Inducible ablation of HUWE1 in hepatocytes suppresses mTORC1 activity in the adult murine liver tissues under fasted and fed conditions.** AAV-TBG-Cre or AAV-TBG-GFP (control) was administered into *Huwe1^flox/y^* male mice (12-week-old) via tail veins (C). AAV-TBG-Cre was administered into control (*Huwe1^+/y^*) or *Huwe1^flox/y^* mice (D). Levels of mTORC1 activity and the indicated protein expression in the liver tissues were monitored by western blotting.

Since Alb-Cre has been shown to start expressing in fetal liver tissues, and the recombination continues during the developmental stage of liver tissues ^51^, we also examined the effect of acute HUWE1 ablation on mTORC1 activity in adult liver tissues (12-week-old) using the adeno-associated viral-thyroxine binding globulin promoter-Cre recombinase (AAV-TBG-Cre) system, which expresses Cre recombinase specifically in the hepatocytes ^52^. Firstly, *Huwe1^f/y^* mice were injected with AAV-TBG-Cre or AAV-TBG-GFP as a control (Fig. 5C). Secondly, the *Huwe1^f/y^* or control wild-type *Huwe1^+/y^*mice were injected with the AAV-TBG-Cre (Fig. 5D). After ten days of infection, mice were sacrificed under fasted or fed conditions, and levels of mTORC1 activity in the liver or kidney tissues were monitored. The expression of HUWE1 was effectively and specifically reduced in only the liver tissues of *Huwe1^f/y^*mice treated with AAV-TBG-Cre (Fig. 5C, 5D, S3B, and S3C). Consistent with the observations in the Alb-Cre system (Fig. 5A and 5B), mTORC1 activity in the liver tissues monitored by levels of phosphorylated S6K1 and 4EBP1 was reduced under both fasted and fed conditions in mice treated with the AAV-TBG-Cre but not with AAV-TBG-GFP (Fig. 5C). Similarly, levels of mTORC1 activity in the liver tissues of *Huwe1^f/y^* mice treated with AAV-TBG-Cre were lower than those of control *Huwe1^+/y^* mice treated with the same AAV-TBG-Cre (Fig. 5D). Together these observations indicate that HUWE1 plays a key role in supporting mTORC1 activity in liver tissues in response to food intake.

### HUWE1 stimulates CAD gene expression and maintains levels of CAD protein in liver tissues

The ablation of HUWE1 slightly but consistently decreased CAD expression in cultured cells (Fig. 4A, 4B, 4D, and S2B). To examine if HUWE1 is also required for maintaining physiological levels of CAD protein expression in murine liver tissues, we monitored CAD expression in *Huwe1 LKO* mice. Importantly, CAD expression was drastically reduced in the liver tissues of *Huwe1 LKO* mice compared to that in *Huwe1^f/y^*control mice (Fig. 6A). However, expression levels of other enzymes in the de novo pyrimidine synthesis pathway (Fig. 6B), including DHODH or UMPS in *Huwe1 LKO* liver remain unchanged (Fig. 6C). Significant reduction of CAD protein expression was also observed in the liver tissues, of which *Huwe1* gene was knocked out in the adult mice by the AAV-TBG-Cre (Fig. 5C and 5D).

**Figure 6.**
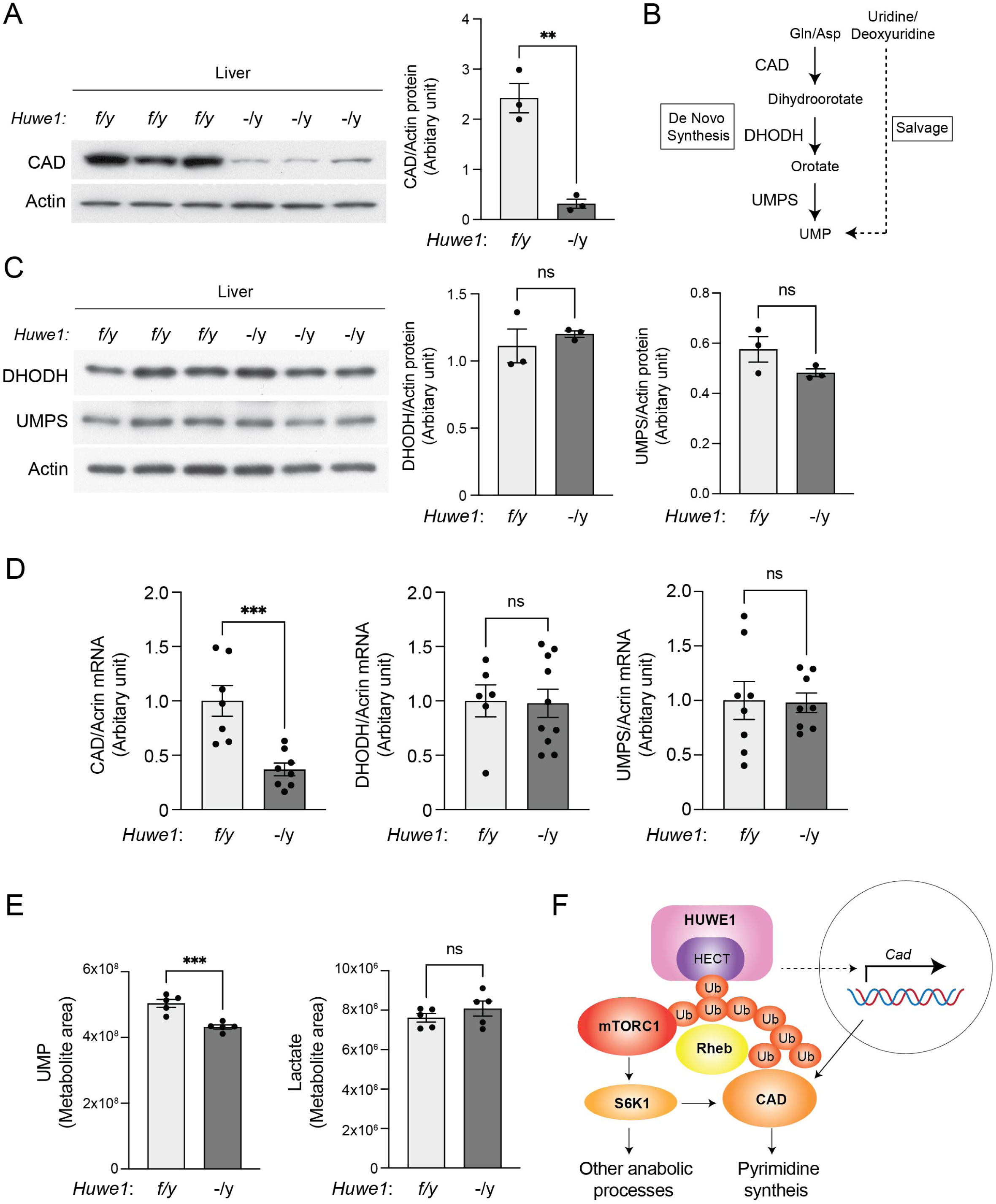
HUWE1 supports de novo pyrimidine synthesis by maintaining CAD expression. **(A) HUWE1 is required for maintaining CAD protein expression in liver tissues.** Levels of CAD protein expression were determined in *Huwe1 LKO* and control (*Huwe1^flox/y^*) liver tissues. Experiments were performed as Fig. 5A. Quantification of CAD protein was expressed as a ratio of CAD/Actin, **p<0.01, mean±SEM, n=3. **(B) Scheme of de novo and salvage pyrimidine synthesis pathways**. **(C) HUWE1 KO does not affect levels of DHODH and UMPS in the liver tissues.** Levels of DHODH and UMPS protein expression were determined in *Huwe1 LKO* and control (*Huwe1^flox/y^*) liver tissues. Experiments were performed as Fig. 5A. Quantification of the proteins was expressed as a ratio of DHODH/Actin or UMPS/Actin, ns (not significant), mean±SEM, n=3. **(D) HUWE1 KO specifically reduces levels of CAD mRNA in the liver tissues.** Levels of CAD, DHODH, and UMPS mRNA were determined in the liver tissues of the indicated mice. Data were expressed as a ratio of the indicated mRNA/actin, ***p<0.001, ns (not significant), mean±SEM, n=6∼10. **(E) HUWE1 KO reduced levels of UMP in the liver tissues**. Levels of UMP and Lactate were determined in the *Huwe1 LKO* and control liver tissues by mass spectrometry analyses. ***p<0.001, ns (not significant), mean±SEM, n=5. **(F) Hypothetical model for HUWE1-dependent regulation of mTORC1 activity and de novo pyrimidine synthesis.** Both HUWE1 and CAD preferentially interact with ubiquitinated Rheb. HUWE1-dependent modification of Ub-Rheb may be required for Ub-Rheb to interact with both CAD and mTOR. Through these processes, HUWE1 plays key roles in supporting Rheb-mTORC1 interaction and subsequent mTORC1-S6K1 activation, leading to CAD activation. Simultaneously, HUWE1 enhances CAD mRNA expression and maintains CAD expression and de novo pyrimidine synthesis.

It has been reported that HUWE1 ubiquitinates c-MYC itself or ubiquitinates and degrades MIZ1, a suppressor of c-MYC, thereby enhancing c-MYC-dependent gene expression ^25,53^. Furthermore, the transcription of the CAD gene is positively regulated by c-MYC ^54,55^. To test if HUWE1 positively regulates CAD expression through its transcriptional activity, levels of CAD mRNA were monitored in HUWE1 KD HEK293T cells. As expected, HUWE1 KD reduced levels of not only CAD mRNA but also other c-MYC-target genes such as CDK6, PPIE, and SLC16A1 (Fig. S3D). Critically, levels of CAD mRNA but not mRNAs encoding other enzymes (i.e., DHODH and UMPS) in the de novo pyrimidine pathway were significantly reduced in *Huwe1 LKO* liver tissues compared to those of control *Huwe1^f/y^*liver tissues (Fig. 6D). These observations indicate that HUWE1 also plays an important role in maintaining CAD protein expression by enhancing *Cad* transcription likely through the activation of c-Myc. Furthermore, while we could not detect measurable levels of dihydroorotate, a final metabolite that is generated by CAD, levels of UMP concentrations were slightly but consistently lower in the liver tissues of *Huwe1 LKO* mice compared to control mice (Fig. 6E). Levels of lactate in the liver tissues, a metabolite indicating glycolysis status and hepatocyte functions (e.g., uptake and clearance) were not changed in *Huwe1 LKO* liver tissues.

Taken together, the data presented in this study propose the model where the HUWE1 functions as a key modulator of Ub-Rheb and acts as a mTORC1 activator by facilitating the interaction between mTORC1 and Rheb. Furthermore, our data suggest that HUWE1 may also function as a key player in supporting de novo pyrimidine synthesis pathway by recruiting CAD into the Rheb-mTORC1 complex and by transcriptionally enhancing CAD gene expression (Fig. 6F).

## Discussion

In this study, we identified HUWE1 as a novel positive regulator of mTORC1 activity. HUWE1 plays a key role in the association of Rheb with mTOR and CAD, a known substrate of Rheb and the mTORC1-S6K1 signaling. Furthermore, HUWE1 is required to maintain levels of CAD expression in both in vitro and in vivo. The ablation of HUWE1 specifically inhibited growth factor or amino acid-induced mTORC1 activation without blocking Akt phosphorylation, Akt-induced TSC2 phosphorylation, the dissociation of TSC2 from the lysosomal membrane, or the recruitment of mTORC1 to the lysosome. In fact, these observations support the idea that HUWE1 functions to activate mTORC1 by facilitating the interaction between Rheb and mTORC1 on the lysosome.

Given that HUWE1 shows a strong binding preference for inactive Rheb species, which are highly ubiquitinated when overexpressed, and display its positive role in activating mTORC1 activity, we suspected that HUWE1 may have a GEF-like activity toward Rheb. However, we failed to detect any nucleotide exchange activity of HUWE1 for Rheb in vitro (Fig. S2A), and rather, we found that the E3 ligase activity of HUWE1 is required for HUWE1-induced mTORC1 activation. Our previous studies indicated that the CUL3-Rbx1-KLHL9 ubiquitin ligase polyubiquitinates Rheb and supports its interaction with mTORC1 and subsequent mTORC1 activation on the lysosome ^20,21^. Ablation of each component of the CUL3-Rbx1-KLHL9 or ectopic expression of ATXN3, a Rheb deubiquitinase, mitigates levels of polyubiquitinated Rheb and mTORC1 activity in cultured cells. Moreover, Rheb lacking four lysine residues (K109/135/151/178) important for the polyubiquitination of Rheb displayed an impairment of its polyubiquitination, association with non-ubiquitinated Rheb and mTORC1, and the ability to enhance mTORC1 activity in response to amino acids and growth factors ^20,21^. HUWE1 is known to ubiquitinate multiple proteins such as N-Myc, Mcl-1, beta-catenin, disheveled, and p53 to regulate cell cycle, growth, proliferation, differentiation, and survival ^25–29,56–58^. HUWE1-mediated protein ubiquitination contributes to not only destabilizing target proteins through its K48-linked ubiquitination but also amplifying ubiquitin-chain-mediated signal transduction. For instance, HUWE1 can activate c-Myc by catalyzing the assembly of K63-linked polyubiquitination chains and inducing the transcriptional activation of a subset of c-Myc target genes ^25^. In addition, HUWE1 also generates K48-linked branched ubiquitin chains on the K63 ubiquitin chains of TRAF6, leading to the amplification of the NF-κB signaling ^24^. We posit that HUWE1 may act as a ubiquitin modifier of polyubiquitinated Rheb by likely making branched chains that may play an important role in amplifying the interaction of Rheb and mTORC1 and subsequent mTORC1 activation. While further studies for the detailed characterization of specific ubiquitin chains of Rheb warrant future investigations, the observations that 1) HUWE1 preferentially interacts with ubiquitinated Rheb, 2) HUWE1 enhances the interaction between mTOR and Rheb, 3) HUWE1 does not influence lysosomal localization of TSC2 as well as mTORC1, 4) HUWE1 does not increase Akt and TSC2 phosphorylation to enhance mTORC1 activity, and 4) HUWE1 increases levels of Rheb ubiquitination, support the idea that HUWE1 may directly act on Rheb to exert its positive role in enhancing cellular mTORC1 activity. Interestingly, a recent study identified that Rheb also receives a neddylation, a ubiquitin-like NEDD8 conjugation, through the UBE2F-SAG ligase on the same lysine residue that is subjected to ubiquitination, and the neddylation of Rheb positively regulate its function to stimulate mTORC1 activity ^20,59^. These new observations also raise the possibility that in addition to ubiquitin, HUWE1, which also acts as an E3 ligase for NEDD8 ^31^, may transfer NEDD8 to the Rheb to confer Ub-Rheb resistance to immediate proteasome-mediated degradation thereby supporting the interaction between Rheb and mTORC1 on the lysosome.

CAD is a known substrate of S6K1, and its activity is enhanced by S6K1-dependent phosphorylation ^33,34^. A recent study postulated that GTP-bound active Rheb interacts with CAD and stimulates the CPS activity of CAD in a manner independent of mTORC1 activity. Consistent with the observations by Sato et al. ^35^, we also found that Rheb preferentially interacts with the CPSase domain of CAD. However, contrary to the model proposed in the previous paper, our observations suggest that the binding of Rheb to CAD is rather Rheb’s ubiquitination dependent regardless of its nucleotide-binding state. In support of this, the ablation of HUWE1 diminished the interaction of CAD with both wild-type and the highly ubiquitinated nucleotide-free Rheb mutant (Fig. 3A and 3B). Given the critical role of mTORC1-S6K1 activity in stimulating CAD activity, it is conceivable that the formation of a multi-protein complex, including mTORC1 and CAD, induced by HUWE1-dependent Rheb ubiquitination would be beneficial for amplifying mTORC1 signaling and expediting the transduction of mTORC1 signal to the de novo pyrimidine synthesis pathway.

Interestingly, we observed that ablation of HUWE1 also reduced CAD protein expression in cultured cells and tissues and found that HUWE1 also plays a key role in stimulating the transcription of CAD. Thus, HUWE1 stimulates not only the enzymatic activity of CAD through the activation of the mTORC1-S6K1 pathway but also the gene expression of CAD. The mechanism by which HUWE1 enhances CAD gene expression remains elusive. However, transcriptional upregulation of the CAD gene, which has the E-box sequence in the promoter, is known to be driven by c-Myc transcription factor ^55,60^. While HUWE1 destabilizes and inhibits N-Myc by catalyzing the assembly of K48-linked polyubiquitination chains ^28,61^, it has been reported that HUWE1-dependent ubiquitination of c-Myc or Miz1, a suppressive component of c-Myc/MAX complex, enhances the E-box associated c-Myc-target gene expression ^25,53^. In fact, ablation of HUWE1 in HEK293T cells suppressed levels of c-Myc-dependent transcripts bearing the E box in their promoter (Fig. S3D). Thus, HUWE1 may also play a key role in maintaining cellular de novo pyrimidine synthesis by enhancing CAD expression in addition to activating its enzymatic activity through mTORC1 activation.

Importantly, in the liver hepatocytes, HUWE1 has been shown to inhibit autophagy by degrading WIPI2, a key protein that is required for autophagosome formation ^30,62^. Interestingly, the study identified that mTORC1-dependent WIPI2 phosphorylation triggers HUWE1-dependent WIPI2 degradation ^63^, demonstrating a cooperative role of mTORC1 and HUWE1 in suppressing autophagosome formation. However, the observations in this study suggest that HUWE1 may exert a broader role in inhibiting the processes of autophagy by supporting mTORC1 activity, which attenuates autophagy in several steps ^64–67^. Moreover, it has recently been reported that mice lacking HUWE1 in their hepatocytes (*Huwe1 LKO*) showed a significant reduction of lipid biogenesis by inhibiting lipogenic proteins and their mRNA expression, such as FASN and ACACB, and displayed resistance to age- and high-fat diet-induced hepatic steatosis ^68^. Interestingly, a recent study discovered that UTP, a key metabolite generated from UMP, plays a key role in supporting pyruvate dehydrogenase activity and subsequent acetyl-CoA synthesis and fatty acid biogenesis, indicative of another key role of pyrimidine synthesis in lipid biogenesis ^69^. Since the mTORC1-S6K1 pathway stimulates both lipid and pyrimidine biogenesis pathways through the activation of SERBPs and CAD, respectively ^33,34,70,71^, it is conceivable that the reduction of mTORC1 activity and CAD expression in *Huwe1 LKO* liver tissues may contribute to protecting aberrant lipid biogenesis and diet-induced non-alcoholic hepatic steatosis.

## Acknowledgment

Metabolomics measurements were performed by the University of Michigan Metabolomics Core (University of Michigan Medical School, Biomedical Research Core Facilities, Ann Arbor, MI, USA). This study was supported by grants from the NIH (DK124709 and GM145631), the DOD (TS140055), and the Takeda Science Foundation.

## Author contributions

T.I., S.H., A.K., S.Y., S.P.S., and Y.Y. performed experiments. H.K. performed AAV-TBG viral infection experiments. H.B. and M.K. measured metabolites. V.B. performed mass spectrometry analyses. A.J. contributed to immunofluorescence analyses. V.S., S.K., A.T., O.K.G., O.K., and F.P. generated and maintained mouse colonies and tissue samples. H.Y. and J.L. helped with data analyses and discussions. T.I. and K.I. conceived the study and experimental design and wrote the manuscript.

## KEY RESOURCES TABLE

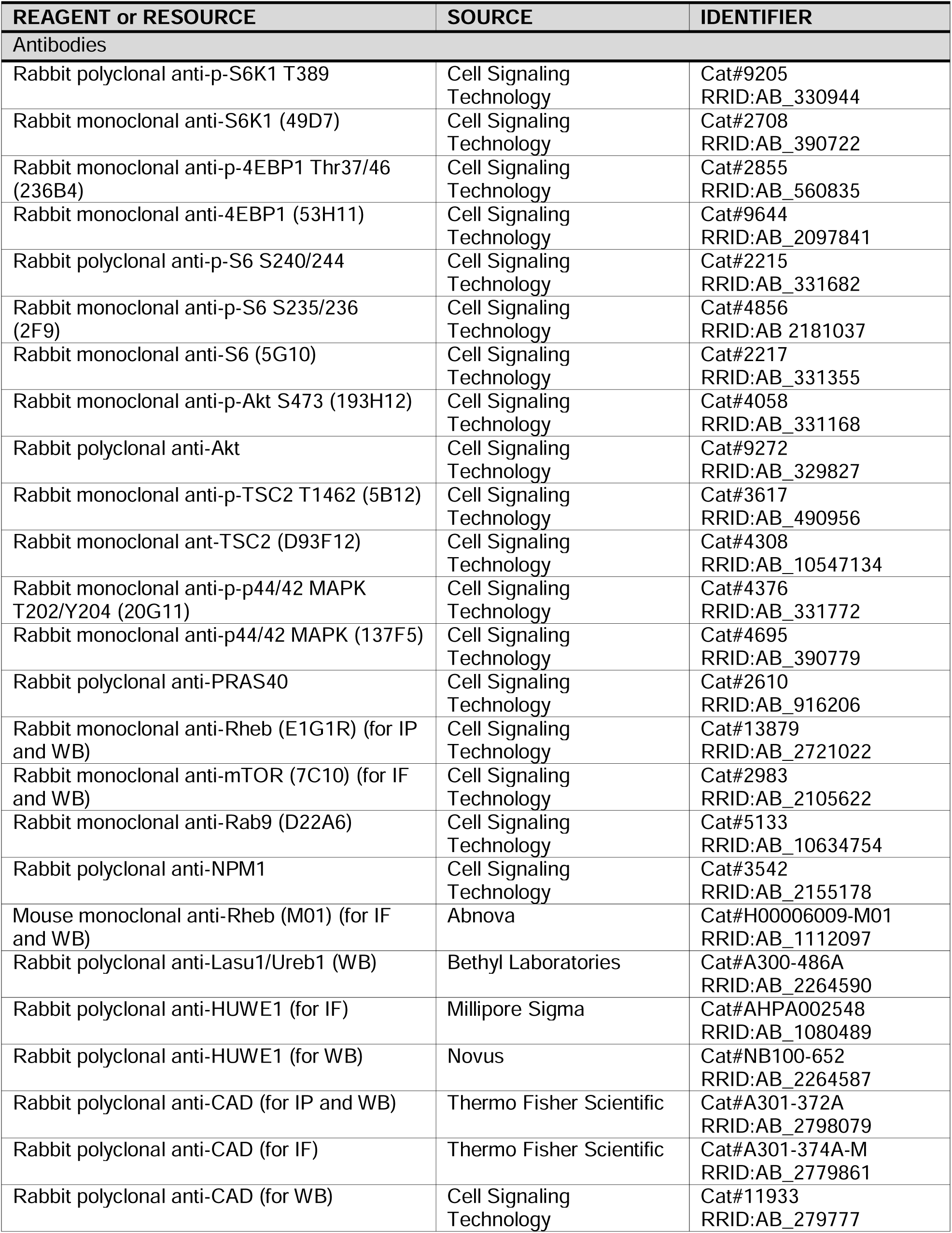
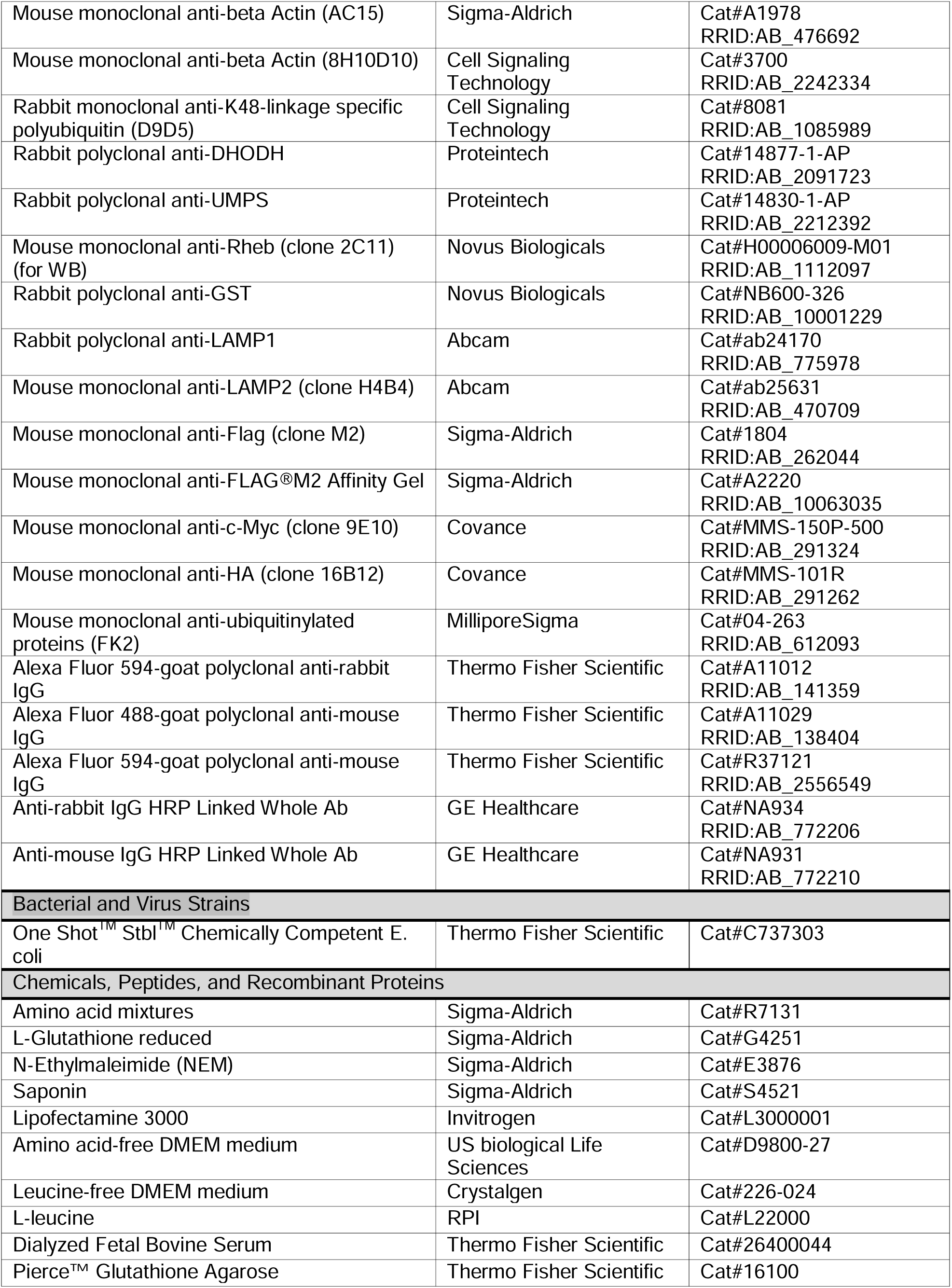
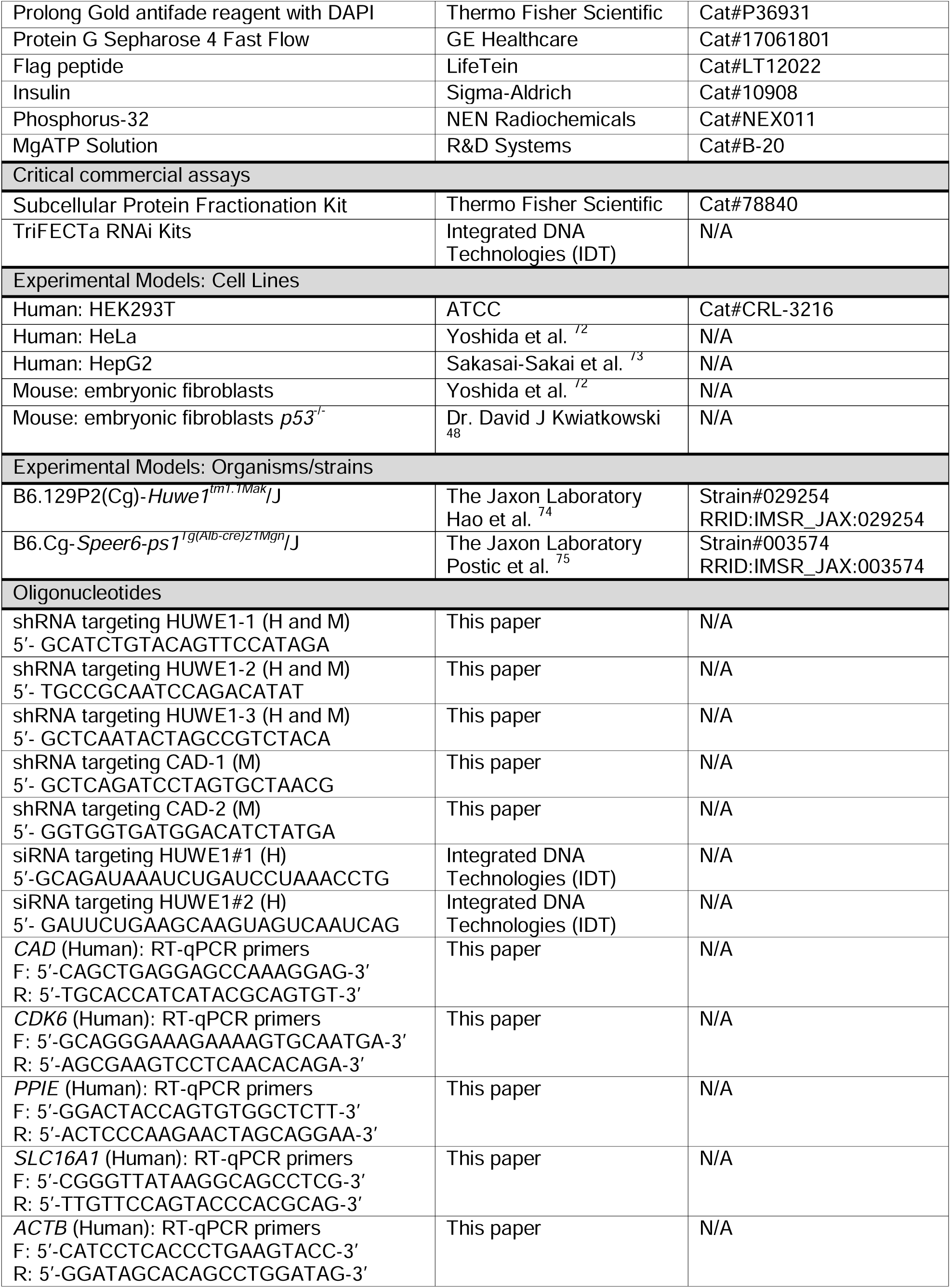
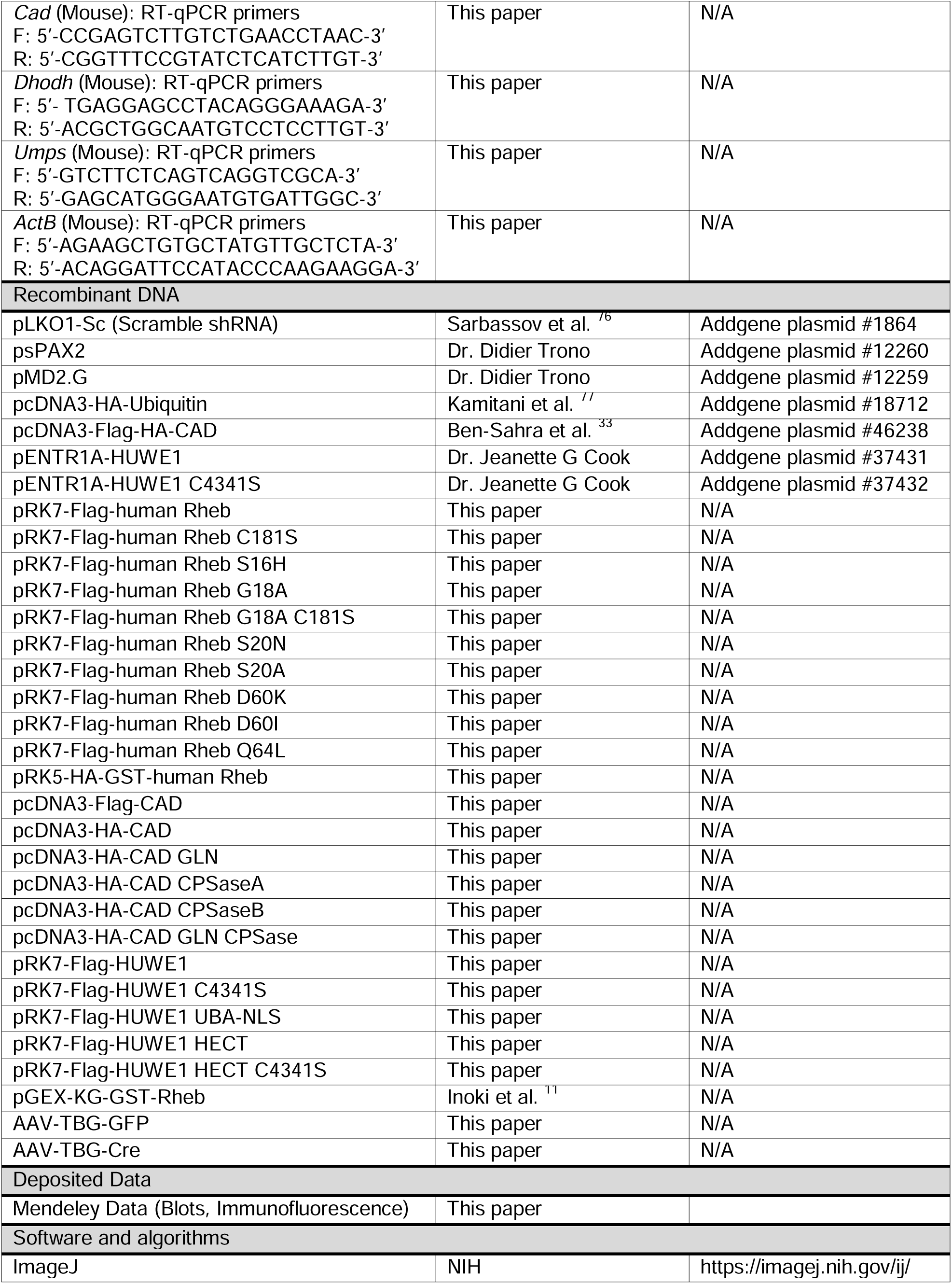
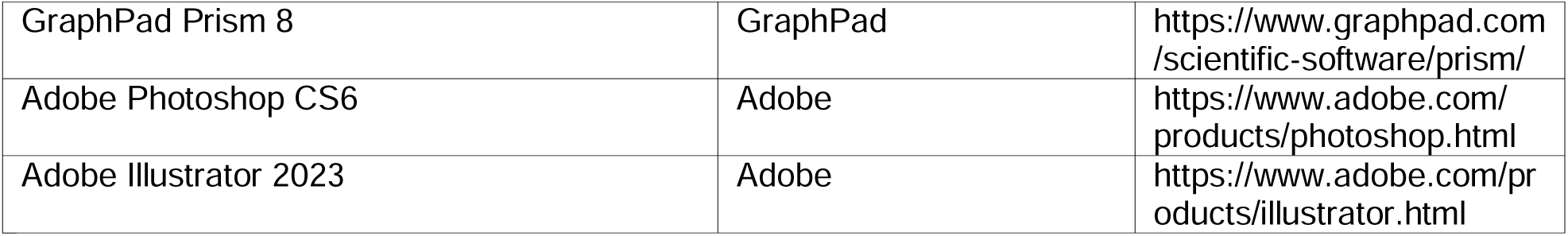

## EXPRIMENTAL MODEL AND STUDY PARTICIPANT DETAILS

### Mouse model

*Huwe1^flox/flox^* (B6) and *Albumin Cre* (B6) mice were obtained from the Jaxon Laboratory^58^. Congenital male HUWE1 liver-specific KO mice (*Huwe1^flox/y^*, *Alb-Cre*) were generated by crossing male *Albumin Cre* (B6) and female *Huwe1^flox/flox^* mice (*Huwe1* is a gene located on the X chromosome). Inducible male HUWE1 liver-specific KO mice were generated by infecting male *Huwe1^flox/y^* mice (12-week-old) with AAV-TBG-Cre virus per tail vein. In analyses of AAV-TBG-Cre-induced HUWE1 KO mice, after ten days infection of control (AAV-TGB-GFP) and AAV-TBG-Cre, mice were starved for 16 hr and fed for 5 hr. The liver and kidney cortex tissues were homogenized with Triton X-100 lysis buffer (50 mM Tris-HCl [pH 7.5], 150 mM NaCl, 1% Triton X-100, 10% glycerol, 10 mM NaF, 10 mM pyrophosphate, 10 mM glycerophosphate, and a mixture of protease inhibitor (Roche)). All animal experiments were approved by the Institutional Animal Care and Research Advisory Committee at the University of Michigan.

### Mammalian Cell lines and treatment

HEK293T, HeLa, HepG2, wild type mouse embryonic fibroblasts (MEFs), and p53-/-MEFs (gifts from Dr. David Kwiatkowski, Brigham and Women’s Hospital Boston, MA, USA) were cultured in DMEM (Invitrogen) with 10% fetal bovine serum (Corning) and 100 units/ml penicillin and 100 µg/ml streptomycin (Invitrogen). All cell lines used in the study were regularly checked for mycoplasma.

## METHOD DETAILS

### Cell culture treatment, transfection, and fractionation

For growth factor/nutrient starvation, cells were incubated with HBSS (Hank’s Balanced Salt Solution, Invitrogen) for the indicated times. For growth factor or nutrients starvation, cells were incubated with FBS-free or leucine (Leu)-free media (Crystalgen) for the indicated time, respectively. For stimulation of these, growth media, insulin (Sigma-Aldrich), or Leu (RPI Corp.) were added for the indicated times. Transfection was performed using Lipofectamine (Invitrogen). MEFs were fractionated to the cytoplasmic, membrane and nuclear fractions using Subcellular Protein Fractionation Kit for Cultured Cells (Pierce).

### shRNAs, lentivirus production

shRNAs were cloned into the lentivirus plasmid pLKO.1-puro. HEK293T cells were transfected with pLKO.1-puro cloned with shRNAs, psPAX2, and pMD2. Media containing lentivirus were collected and spun at 26,000 rpm for 100 min. Virus pellets were resuspended with Opit-MEM (Invitrogen) and stored at -80°C until use. Cells were infected for 24 hr and selected with 2 µg/ml puromycin. 72 hr post-infection, cells were harvested and processed for further analysis.

### siRNA experiments

TriFECTa RNAi Kits were purchased from Integrated DNA Technologies (IDT). Two sequences for human HUWE1 were used: GCAGAUAAAUCUGAUCCUAAACCTG (hs.Ri.HUWE1.13.1) and GAUUCUGAAGCAAGUAGUCAAUCAG (hs.Ri.HUWE1.13.2). Non-targeting DsiRNA included in the kit was used as a negative control. siRNAs were pre-mixed with RNAiMAX (Invitrogen) for 10 min. HeLa cell suspension was seeded onto 6-well plate at 2-3 x 10^5^ cells/2.5 mL. siRNA/RNAiMAX mixture was added to the cells, and then cells were incubated for 48 hr.

### Cell lysis and immunoprecipitation

Cells were harvested with NP40 lysis buffer (10 mM Tris-HCl [pH 7.5], 2 mM EDTA, 100 mM NaCl, 1% NP-40, 60 mM NaF, 10 mM pyrophosphate, 10 mM glycerophosphate, and a mixture of protease inhibitor (Roche)) for western blot analyses, CHAPS lysis buffer (40 mM HEPES [pH 7.5], 120 mM NaCl, 1 mM EDTA, 0.3% CHAPS, 50 mM NaF, 10 mM pyrophosphate, 30 mM glycerophosphate, and a mixture of protease inhibitor) for immunoprecipitation to assess protein-protein interaction, or RIPA lysis buffer (25 mM Tris-HCl [pH7.6], 50 mM NaCl, 1% NP-40, 1% sodium deoxycholate, 0.1% SDS, 50 mM NaF, 10 mM pyrophosphate, 10 mM glycerophosphate, and a mixture of protease inhibitor) for in vivo crosslinking samples. For immunoprecipitation using Flag-agarose beads, 15 µl of anti-FLAG M2 affinity gel (Sigma-Aldrich) was added and incubated with gentle rocking for 3 hr at 4°C. For immunoprecipitation using the specific antibodies, 1 µg of antibody was added and incubated with gently rocking for 3 hr at 4°C. 15 µl of 50% slurry of protein G sepharose (GE Healthcare) was added and incubated for additional 1 hr. Immunoprecipitates were washed with lysis buffer for five times and then denatured at 100°C for 5 min in 1X SDS sample buffer.

### Immunofluorescence staining of cultured cells

MEFs were plated on glass coverslips in a 6-well plate. Coverslips were rinsed with PBS and fixed with 4% paraformaldehyde (PFA) for 15 min and then permeabilized with 0.1% saponin and 2% BSA in PBS. After three washes with PBS, coverslips were incubated with the indicated primary antibodies in PBS containing 0.1% saponin and 2% BSA overnight at 4°C. The coverslips were washed 3 times with PBS and incubated with secondary antibodies (1:500) in 0.1% saponin/2% BSA for 2 hr at room temperature. After 3 washes with PBS, the coverslips were mounted on the glass slides using Prolong Gold antifade reagent with DAPI (Invitrogen) and analyzed with a Leica TCS SP5 confocal microscope. The colocalization ratio was measured using the Fiji package of ImageJ (https://fiji.sc). Briefly, a binary image was created by Otsu’s Threshold algorithm. The overlapped pixels were obtained by merging two binary images. The total pixel particles of each binary image and the merged image were measured by Analyze Particles. The ratio was calculated by the endogenous Rheb co-localized with endogenous HUWE1 or CAD to total endogenous Rheb, and the HUWE1 or CAD co-localized with Rheb to total HUWE1 or CAD.

### Immunofluorescence staining of tissues and measurements of signals

The liver tissues (right posterior segment VI) were fixed with PBS containing 4% PFA. The fixed tissues were embedded into paraffin and sectioned in 5 µm and subjected to immunofluorescence staining. Antibodies were used at the following dilutions: pS6 (1:400; Cell Signaling Technology); pAkt S473 (1:400; Cell Signaling Technology). Prolong Gold antifade reagent with DAPI (Invitrogen) was used for nucleus staining. The number of hepatocytes and the intensity of fluorescence signals were measured by ImageJ software.

Briefly, the area was calculated with a standard measurement bar of 20 µm in each image. Vascular regions were eliminated from the area. Relative TRITC intensity (pS6 and pAkt) in the area was determined through ImageJ’s measurement analysis tool. The DAPI channel was used to determine the number of hepatocyte nuclei by counting the particles of an area larger than 20 µm^2^.

### RT-qPCR

Cells were collected and lysed with TRIzol reagent (Invitrogen). Total RNA was purified using the RNeasy Mini Kit (QIAGEN). RT-qPCR was performed to determine the relative mRNA levels of human CAD, CDK6, PPIE, and SLC16A1 and murine CAD, DHODH, and UMPS using gene-specific primers (Supplemental information), the One Step TB Green PrimeScript PLUS RT-PCR kit (Takara), and the StepOnePlus Real-time PCR System (Applied Biosystems).

### In vivo labeling

GTP- and GDP-bound Rheb was determined by in vivo labeling with phosphorus-32 (^32^P) radionucleotide as described previously ^11,40^. Briefly, cells were washed once and incubated with phosphate-free DMEM (Invitrogen) for 90 min and incubated with phosphate-free DMEM containing 10% dialyzed FBS and 25µCi ^32^P-phosphate/ml for 4 hr. Cells were lysed with cell lysis buffer (0.5% NP-40, 50 mM Tris-HCl [pH 7.5], 100 mM NaCl, 10 mM MgCl_2_, 1 mM DTT, and a mixture of protease inhibitor) at 4°C for 30 sec. The lysate was then centrifuged at 16,000 x g for 15 min at 4°C. To immunoprecipitate Flag-tagged Rheb, 10 µl of anti-FLAG M2 affinity gel was added to the supernatant and incubated with gentle rocking for 2 hr at 4°C. The beads were washed with wash buffer 1 (50 mM Tris-HCl [pH 8.0], 500 mM NaCl, 5 mM MgCl_2_, 1 mM DTT, 0.5% Triton X-100) three times at 4°C and then with wash buffer 2 (50 mM Tris-HCl [pH 8.0], 100 mM NaCl, 5 mM MgCl_2_, 1 mM DTT, 0.1% Triton X-100) three times at 4°C. GTP and GDP nucleotides bound to Rheb were eluted with the elution buffer (2 mM EDTA, 0.2% SDS, 1 mM GDP, and 1 mM GTP) for 10 min at 68°C. Aliquots were spotted on PEI cellulose plates (Millipore) and resolved by thin-layer chromatography (TLC) with 0.75 M KH_2_PO_4_ (pH 3.4). Radionucleotides were detected with PhosphorImager (GE Healthcare) and quantified by Image J. The following formula is used to calculate the percentage of GTP: %GTP = (GTP/[(GDP x 1.5) + GTP]) x 100.

### Protein identification by LC-Tandem MS

In-gel digestion: Immunoprecipitants with the indicated Flag-Rheb mutants were separated by SDS-PAGE and stained with Coomassie Blue G-250 (ThermoFisher Scientific). Gel slice, including the specific bands co-IPed with Flag-Rheb G18A mutant, was destained with 30% methanol for 4 h. Upon reduction (10 mM DTT) and alklylation (65 mM 2-Chloroacetamide) of the cysteines, proteins were digested overnight with 500 ng of sequencing grade, modified trypsin (Promega) at 37° C. Peptides were extracted by incubating the gel with 150 µl of 50% acetonitrile/0.1% TFA for 30 min at room temperature. A second extraction with 150 µl of 100% acetonitrile/0.1% TFA was also performed. Both extracts were combined and dried in a vacufuge.

Liquid Chromatography and tandem mass spectrometry (LC-MS/MS): The resulting peptides were dissolved in 9 µl (for in-gel digest) or an appropriate volume of 0.1% formic acid/2% acetonitrile solution (to achieve ∼500ng peptide/ul), and 2 µls of the peptide solution were resolved on a nano-capillary reverse phase column (Acclaim PepMap C18, 2 microns, 50 cm, (ThermoFisher Scientific)) using a 0.1% formic acid/2% acetonitrile (Buffer A) and 0.1% formic acid/95% acetonitrile (Buffer B) gradient at 300 nl/min over a period of 90 min (2-25% buffer B in 45 min, 25-40% in 5 min, 40-90% in 5 min followed by holding at 90% buffer B for 5 min and reequilibration with Buffer A for 30 min). Eluent was directly introduced into Orbitrap Fusion tribrid mass spectrometer (ThermoFisher Scientific) using an EasySpray source. MS1 scans were acquired at 120K resolution (AGC target=1x10^6^; max IT=50 ms). Data-dependent collision-induced dissociation MS/MS spectra were acquired using the Top speed method (3 seconds) following each MS1 scan (NCE ∼32%; AGC target 1x10^5^; max IT 45 ms).

Database Search: Proteins were identified by searching the data against UniProt Homo sapiens protein database appended with common contaminant protein list (ftp://ftp.thegpm.org/fasta/cRAP) using Proteome Discoverer (v2.0, ThermoFisher Scientific). Search parameters included MS1 mass tolerance of 10 ppm and fragment tolerance of 0.1 Da; two missed cleavages were allowed; carbamidomethylation of cysteine, oxidation of methionine, and deamidation of asparagine and glutamine, were considered as potential modifications. FixedPSM validator of Proteome Discoverer was used to filter and retain high quality proteins/peptides.

The protein samples were processed and analyzed at the Mass Spectrometry Facility of the Department of Pathology at the University of Michigan.

### UMP and Lactate measurement

Central Carbon Metabolism and Glycolysis/TCA sample preparation: Liver samples were removed from -80 °C storage and maintained on wet ice throughout the processing steps. Samples were carefully resected and weighed to 30 mg +/- 3mg into a microtube containing stainless steel beads for tissue disruption, and the extraction solvent (1:1:1:1= Methanol:Acetonitrile:Acetone:Water) was added at a ratio of 50 µL/mg of tissue. Tissue disruption was performed in a Precellys Evolution Touch homogenizer set to 0°C, 6200 rpm, for 3 cycles (20 sec shake time and 30 sec rest time per cycle.) After removal from the homogenizer, samples were incubated on ice for 10 minutes, vortexed to remix, then centrifuged at 14,000 RPM for 10 min at 4°C. 20 µL of the supernatant from each sample was removed to create a pooled sample for QC purposes. For both the pool and samples, 200 µL of the extract was transferred to an autosampler vial and dried under a gentle steam of nitrogen at room temperature. Dried samples were reconstituted in 100 µL of 80/20 water/methanol for LC-MS analysis. TCA and Central Carbon Metabolism LC-MS analysis: Samples were analyzed on an Ion Pairing LC-MS system consisting of an Agilent Infinity Lab II UPLC coupled with a 6545 QTOF mass spectrometer in negative ion mode, as previously described ^78^. Semi-quantitative data for UMP and Lactate were obtained by manual integration using Profinder v8.00 (Agilent Technologies, Santa Clara, CA.) Metabolites were identified by matching the retention time (+/- 0.1 min), mass (+/- 10 ppm) and isotope profile (peak height and spacing) to authentic standards run at the same time as the samples.

### Statistical analysis

The significance of differences was determined using an unpaired two-tailed Student’s t-test or one-way ANOVA. p values of less than 0.05 were considered statistically significant: *p<0.05, **p<0.01, ***p<0.001, ****p<0.0001.

## Supplemental Figure Legends

**Supplemental Figure 1 (related to Figure 1 and 2)**

**(S1A) HUWE1 and CAD express in the cytosolic and membrane fractions.** MEFs were cultured under steady-state growth (Basal), serum-starved (1 hr, DMEM without FBS), or insulin-stimulated (15 min) conditions and fractionated as described in the Method section. Levels of the indicated proteins were monitored.

**(S1B) Endogenous HUWE1 partially co-localizes with endogenous Rheb.** MEFs expressing control or HUWE1 shRNA were monitored by double immunofluorescence staining with HUWE1 (green) and Rheb (red) antibodies.

**(S1C) Endogenous CAD partially co-localizes with endogenous Rheb.** MEFs expressing control or CAD shRNA were monitored by performing double immunofluorescence staining with CAD (green) and Rheb (red) antibodies.

**(S1D) Quantification of co-localization.** Pearson’s correlation coefficient and R values were determined for quantifying co-localization of HUWE1/Rheb and CAD/Rheb and graphed as a mean ± SEM, n=8.

**(S1E) Levels of HUWE1/Rheb co-localization.** Percentages of HUWE1/Rheb co-localization were analyzed by Otsu’s Threshold Algorism. The graph expressed as a ratio (co-localization of the indicated protein/total indicated protein). Data were expressed as a mean±SEM, n=30.

**(S1F) Levels of CAD/Rheb co-localization.** Percentages of CAD/Rheb co-localization were analyzed by Otsu’s Threshold Algorism. The graph expressed as a ratio (co-localization of the indicated protein/total indicated protein). Data were expressed as a mean±SEM, n=25.

**Supplemental Figure 2 (related to Figure 1 and 4)**

**(S2A) HUWE1 did not display a GEF activity toward Rheb in vitro.** GST-Rheb purified from bacteria was loaded with radioactive GDP (^32^P-GDP) and incubated with GTPγS in the presence or absence of immunopurified HUWE1 from HEK293T cells for the indicated times in vitro. Levels of remaining radioactive GDP with GST-Rheb were measured by scintillation counter. Data were expressed as mean±SEM, n=3. ns (not significant).

**(S2B) HUWE1 KD inhibits mTORC1 activity in HEK293T cells.** HEK293T cells were infected with HUWE1 shRNAs and levels of mTORC1 activity and the indicated protein expression were monitored under steady-state growth culture (Basal), HBSS starved (1 hr, HBSS), or growth factor/amino acid-stimulated (20 min, DMEM containing 10% FBS) conditions.

**(S2C) HUWE1 KD reduces mTORC1 activity in HeLa cells.** HeLa cells were transfected with HUWE1 siRNAs or control siRNA, and levels of mTORC1 activity and the indicated protein expression were monitored under steady-state growth culture condition (Basal).

**(S2D) HUWE1 KD does not influence insulin-induced dissociation of TSC2 from the lysosome.** MEFs knocked down HUWE1 expression were serum-starved then stimulated with insulin (1 µg/ml) and lysosomal localization of TSC2 (red) and LAMP2 (green) was monitored by performing double immunofluorescence staining. Pearson’s correlation coefficient, R values, were determined for quantifying co-localization of TSC2 and LAMP2 and graphed as a mean±SEM (***p<0.001). n=19 (Scr, -Ins), n=27 (Scr, +Ins), n=14 (sh HUWE1, -Ins), n=12 (sh HUWE1, +Ins). ns (not significant).

**Supplemental Figure 3 (related to Figure 5 and 6)**

**(S3A) Congenital ablation of HUWE1 in hapatocytes had little effect on Akt phosphorylation in the murine liver tissues.** *Huwe1^flox/y^* and *Huwe1 LKO* (*Huwe1^flox/y^*, *Alb-Cre*) mice were starved for 16 hr and fed for 5 hr. mTORC1 activity and nucleus number of liver tissues (right posterior segment VI) were monitored by staining with pAkt (S473) antibody and DAPI. Quantification was demonstrated as nucleus number of hepatocyte/area (counts/µm^2^) and the intensity of pAkt/nucleus number of hepatocyte (arbitrary unit). Ns (not significant), mean±SEM, n=16∼20 images.

**(S3B and S3C) The effect of AAV-TBG-Cre on mTORC1 activity in the kidney tissues of HUWE1 flox/y mice.** AAV-TBG-Cre or AAV-TBG-GFP was administered into the indicated *Huwe1^flox/y^*mice (S3B). AA-TBG-Cre was administrated into control (wild-type, *Huwe1^+/y^*) or *Huwe1^flox/y^* mice. Levels of mTORC1 activity (p240/244 S6) and the indicated protein expression in the kidney tissues were monitored. Note that levels of pS6K1 and S6K1 were too low to convincingly be monitored by immunoblots, and levels of CAD protein expression were maintained in the kidney of the mice treated with AA-TBG-Cre.

**(S3D) HUWE1 KD reduced levels of CAD mRNA expression in HEK293T cells.** Levels of CAD mRNA and other c-Myc target transcripts (CDC25, TOP1, CDK6, PPIE, and SLC16A1) were monitored in control and HUWE1 KD HEK293T cells by RT-qPCR. Data were expressed as mean±SEM, ****p<0.0001, n=12.

