## Supplementary material for "HUWE1 stimulates mTORC1 activity by enhancing Rheb interaction with mTORC1 and supports de novo pyrimidine synthesis": online supplemental file: Supplemental Figures and texts.pdf

Supplemental Figure 1

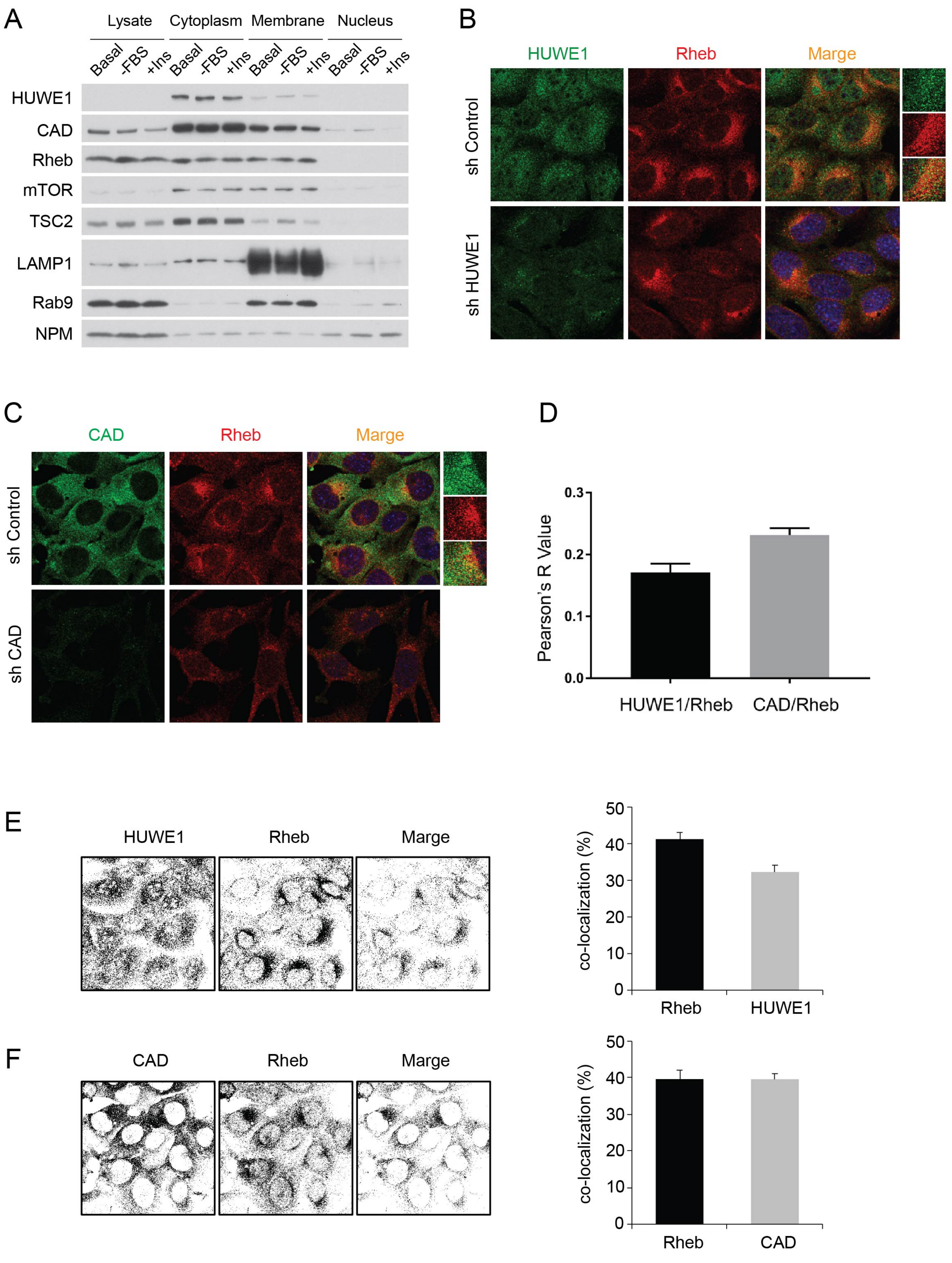

#### Supplemental Figure Legends

##### Supplemental Figure 1 (related to Figure 1 and 2)

**(S1A) HUWE1 and CAD express in the cytosolic and membrane fractions.** MEFs were cultured under steady-state growth (Basal), serum-starved (1 hr, DMEM without FBS), or insulin-stimulated (15 min) conditions and fractionated as described in the Method section. Levels of the indicated proteins were monitored.

**(S1B) Endogenous HUWE1 partially co-localizes with endogenous Rheb.** MEFs expressing control or HUWE1 shRNA were monitored by double immunofluorescence staining with HUWE1 (green) and Rheb (red) antibodies.

**(S1C) Endogenous CAD partially co-localizes with endogenous Rheb.** MEFs expressing control or CAD shRNA were monitored by performing double immunofluorescence staining with CAD (green) and Rheb (red) antibodies.

**(S1D) Quantification of co-localization.** Pearson's correlation coefficient and R values were determined for quantifying co-localization of HUWE1/Rheb and CAD/Rheb and graphed as a mean ± SEM, n=8.

**(S1E) Levels of HUWE1/Rheb co-localization.** Percentages of HUWE1/Rheb co-localization were analyzed by Otsu's Threshold Algorism. The graph expressed as a ratio (co-localization of the indicated protein/total indicated protein). Data were expressed as a mean±SEM, n=30.

**(S1F) Levels of CAD/Rheb co-localization.** Percentages of CAD/Rheb co-localization were analyzed by Otsu's Threshold Algorism. The graph expressed as a ratio (co-localization of the indicated protein/total indicated protein). Data were expressed as a mean±SEM, n=25.

### Supplemental Figure 2

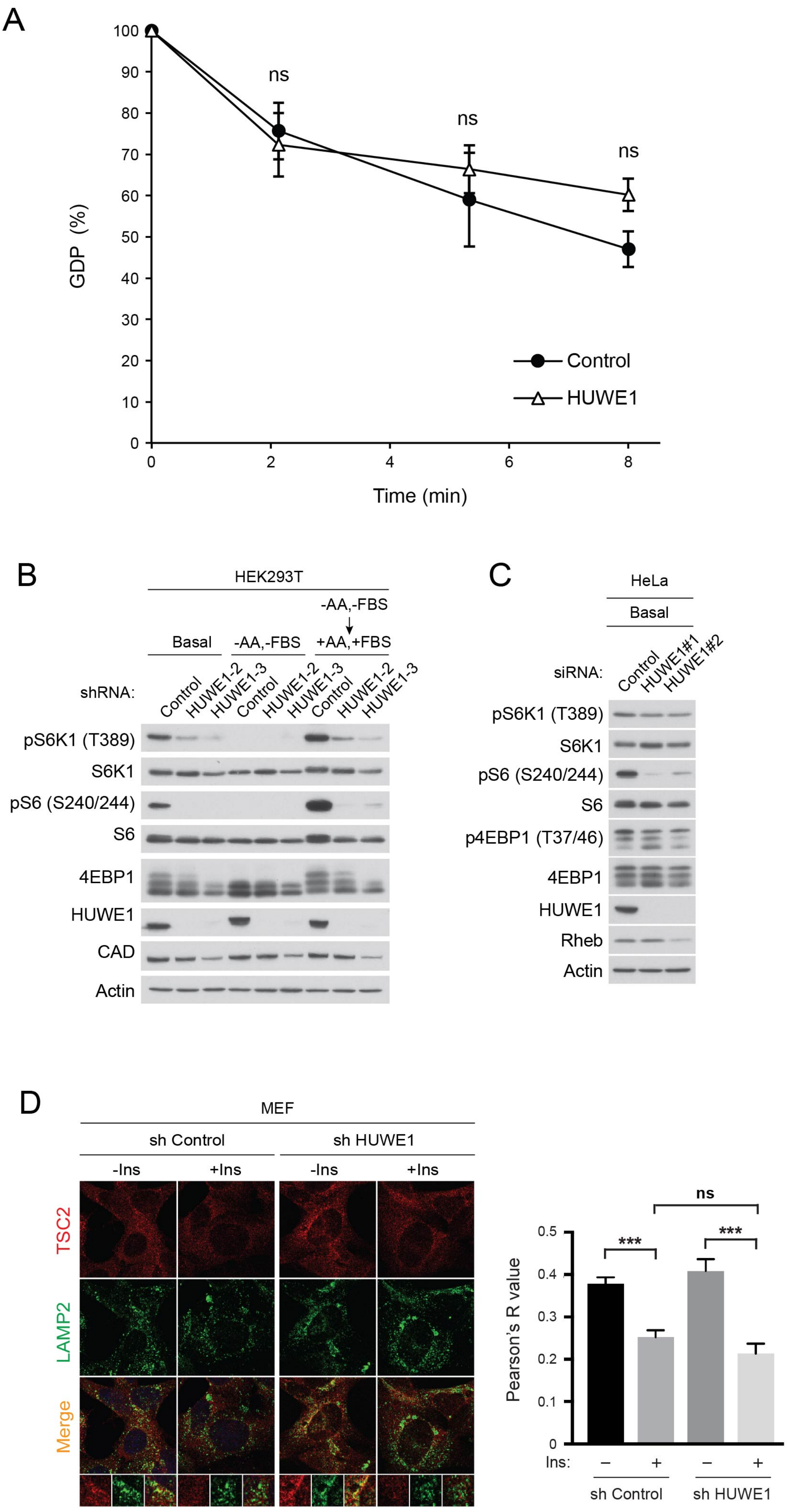

**Supplemental Figure 2 (related to Figure 1 and 4)**

**(S2A) HUWE1 did not display a GEF activity toward Rheb in vitro.** GST-Rheb purified from bacteria was loaded with radioactive GDP ($^{32}$P-GDP) and incubated with GTPγS in the presence or absence of immunopurified HUWE1 from HEK293T cells for the indicated times in vitro. Levels of remaining radioactive GDP with GST-Rheb were measured by scintillation counter. Data were expressed as mean±SEM, n=3. ns (not significant).

**(S2B) HUWE1 KD inhibits mTORC1 activity in HEK293T cells.** HEK293T cells were infected with HUWE1 shRNAs and levels of mTORC1 activity and the indicated protein expression were monitored under steady-state growth culture (Basal), HBSS starved (1 hr, HBSS), or growth factor/amino acid-stimulated (20 min, DMEM containing 10% FBS) conditions.

**(S2C) HUWE1 KD reduces mTORC1 activity in HeLa cells.** HeLa cells were transfected with HUWE1 siRNAs or control siRNA, and levels of mTORC1 activity and the indicated protein expression were monitored under steady-state growth culture condition (Basal).

**(S2D) HUWE1 KD does not influence insulin-induced dissociation of TSC2 from the lysosome.** MEFs knocked down HUWE1 expression were serum-starved then stimulated with insulin (1 μg/ml) and lysosomal localization of TSC2 (red) and LAMP2 (green) was monitored by performing double immunofluorescence staining. Pearson's correlation coefficient, R values, were determined for quantifying co-localization of TSC2 and LAMP2 and graphed as a mean±SEM (***p<0.001). n=19 (Scr, -Ins), n=27 (Scr, +Ins), n=14 (sh HUWE1, -Ins), n=12 (sh HUWE1, +Ins). ns (not significant).

### Supplemental Figure 3

A

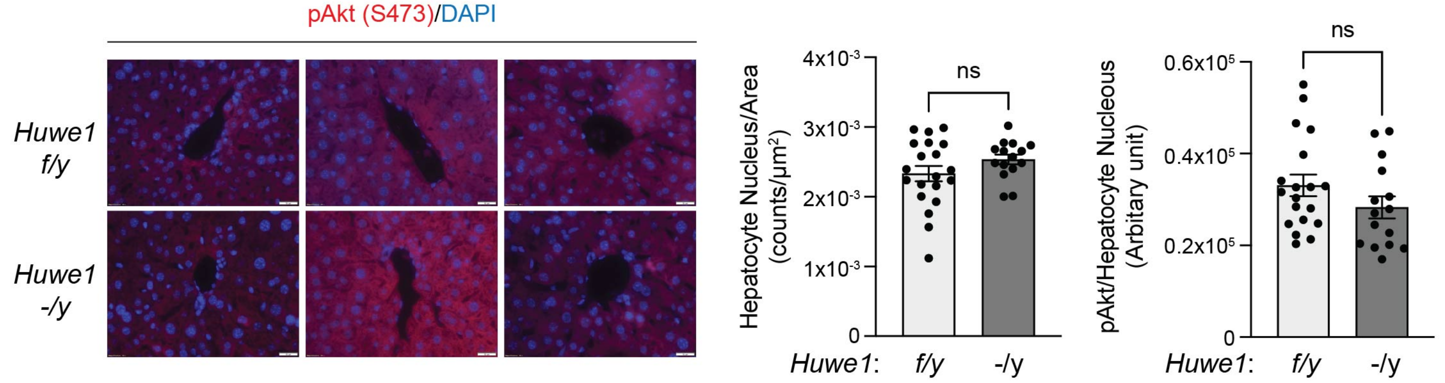


B

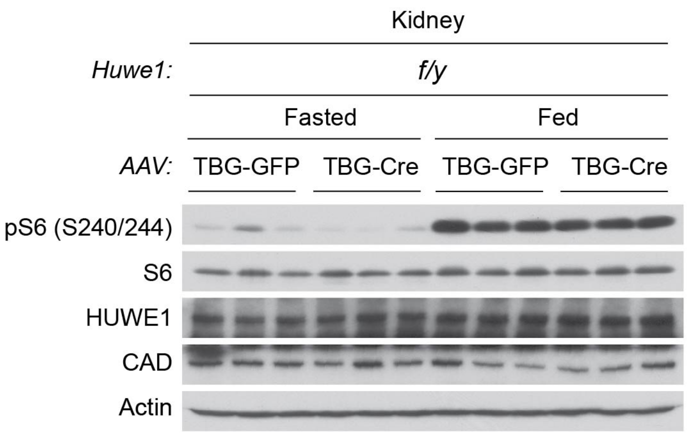


C

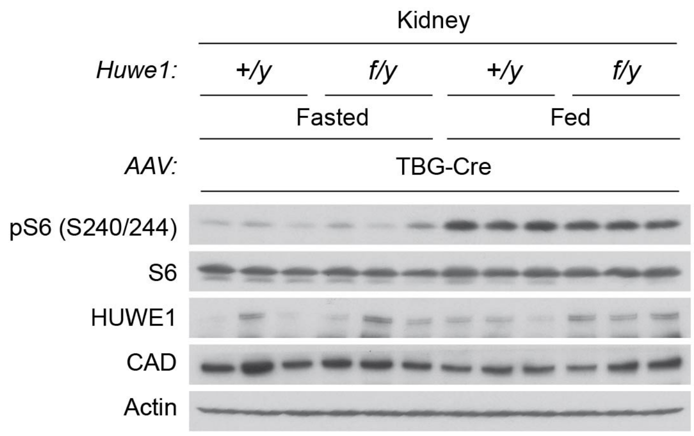


D

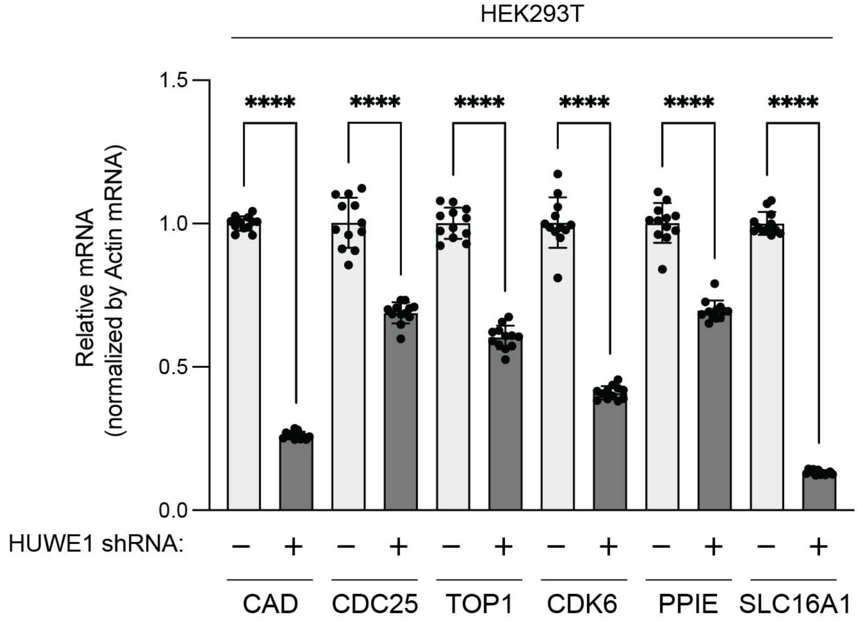

**Supplemental Figure 3 (related to Figure 5 and 6)**

**(S3A) Congenital ablation of HUWE1 in hapatocytes had little effect on Akt phosphorylation in the murine liver tissues.** *Huwe1$^{flox/y}$* and *Huwe1 LKO* (*Huwe1$^{flox/y}$*, *Alb-Cre*) mice were starved for 16 hr and fed for 5 hr. mTORC1 activity and nucleus number of liver tissues (right posterior segment VI) were monitored by staining with pAkt (S473) antibody and DAPI. Quantification was demonstrated as nucleus number of hepatocyte/area (counts/$\mu m^2$) and the intensity of pAkt/nucleus number of hepatocyte (arbitrary unit). Ns (not significant), mean±SEM, n=16~20 images.

**(S3B and S3C) The effect of AAV-TBG-Cre on mTORC1 activity in the kidney tissues of HUWE1 flox/flox mice.** AAV-TBG-Cre or AAV-TBG-GFP was administered into the indicated *Huwe1$^{flox/y}$* mice (S3B). AA-TBG-Cre was administrated into control (wild-type, *Huwe1$^{+/y}$*) or *Huwe1$^{flox/y}$* mice. Levels of mTORC1 activity (p240/244 S6) and the indicated protein expression in the kidney tissues were monitored. Note that levels of pS6K1 and S6K1 were too low to convincingly be monitored by immunoblots, and levels of CAD protein expression were maintained in the kidney of the mice treated with AA-TBG-Cre.

**(S3D) HUWE1 KD reduced levels of CAD mRNA expression in HEK293T cells.**
Levels of CAD mRNA and other c-Myc target transcripts (CDC25, TOP1, CDK6, PPIE, and SLC16A1) were monitored in control and HUWE1 KD HEK293T cells by RT-qPCR. Data were expressed as mean±SEM, ****p<0.0001, n=12.
